# Characterization of dynamic age-dependent changes and driver microbes in primate gut microbiota during host’s development and healthy aging via captive crab-eating macaque model

**DOI:** 10.1101/2020.03.30.015305

**Authors:** Zhi-Yuan Wei, Jun-Hua Rao, Ming-Tian Tang, Guo-An Zhao, Qi-Chun Li, Li-Ming Wu, Shao-Qiang Liu, Bi-Hai Li, Bai-Quan Xiao, Xing-Yin Liu, Jian-Huan Chen

## Abstract

Recent population studies have significantly advanced our understanding of how age shapes the gut microbiota. However, the actual role of age could be inevitably confounded due to varying environmental factors in human populations. A well-controlled environment is thus necessary to reduce undesirable cofounding effects, and recapitulate age-dependent taxonomic and functional changes in the healthy primate gut microbiota. Herein we performed 16S rRNA gene sequencing, characterized age-associated gut microbial profiles from infant to elderly crab-eating macaques reared in captivity, and systemically revealed lifelong dynamic changes of primate gut microbiota in the model. While the most significantly age-associated gut microbial taxa were mainly found in commensals such as *Faecalibacterium*, a set of suspicious pathogens such as *Helicobacter* were exclusively enriched in infants, pointing to their potential role in host development. Importantly, topology analysis indicated that the connectivity of gut microbial network was even more age-dependent than taxonomic diversity, with its tremendous decline probably linked to the host’s healthy aging. NetShift analysis identified *Prevotella 9, Rikenellaceae RC9 gut group* and *Megasphaera* as key drivers during gut microbiota maturation and development, actively involved in age-dependent changes in phenotypes and functions of the gut microbial community. The current study demonstrates lifelong age-dependent changes in healthy primate gut microbiota. Our findings indicate potential importance of appropriate exposure to suspicious pathogens in infant development. The age-associated baseline profiles and driver microbes of primate gut microbiota in the current study could provide new insight into its role in the host’s development and healthy aging.

## Introduction

The human gut microbiota is composed of trillions of microbial cells that habitat in the gastrointestinal tract[1]. These microbes altogether encode an extremely large and dynamic genetic diversity, enabling the host to access additional energy and metabolites [2]. The gut microbiota thus plays a substantial role in human physiology and health [3]. In particular, commensal microbes in the gastrointestinal tract interplay with the host immune system, protect the host from pathogens, and modulate the host’s physiological functions with commensal-derived metabolites [4-6].

The development of human gut microbiota, with dynamic changes after birth, have been implicated to play an active role concomitantly with the host’s development and aging [7]. After first colonization at birth, the postnatal gut microbiota develops rapidly in the first few months of life [8, 9]. By 1□week of age, the infant gut microbiota has already become very similar to that at one-month old [10]. Breastfeeding is one of the key factors that greatly shape the infant gut microbiota, and is linked to the increase of *Bifidobacterium* species [11]. Analysis of fecal bacteria in human populations shows that changes may occur in the gut microbiota as age increases, and could be associated with increased risk of disease, especially age-related diseases such as type 2 diabetes and hypertension in elderly people [7, 12-14].

Nevertheless, the actual effects of age on human gut microbiota remain to be further elucidated. The human gut microbial community is known to be highly dynamic. The existing population-based studies are inevitably influenced by a number of confounding factors in the populations. The individual human microbiota pattern is vastly variable. And varying environmental factors, such as diets [15] and antibiotic use [16] could dramatically influence the bacterial community [17]. In addition, people of different generations in the same population may have distinct growth experience and life styles due to the rapid urbanization of most human societies, which also shape the human gut microbiota [18]. These confounding factors emphasized the difficulty and importance to study healthy core native gut microbiota. A well-controlled model system that faithfully recapitulates age-dependent changes in the gut microbiota is thus needed, and would provide better understanding of the role played by the gut microbiota in the host’s healthy development and aging. In addition, humans have a much longer life span and evident difference in the gut microbiota compared to rodents, the lab animals the most widely used in existing gut microbiome studies[19]. In contrast, non-human primates (NHPs) have high similarities to humans in genetics, physiology as well as gut microbial compositions [20]. Moreover, NHPs in captivity have been found to have physiological characteristics and gut microbiota composition similar to those in humans [21]. Captive NHPs are reared with a formula diet and a stable environment, providing a feasible model to study age-dependent changes in the gut microbiota of humans and NHPs.

Various microbes in the gut microbiota interact to form a complex biological network. Therefore, not only taxonomic compositions, but also microbial interactions are essential to infer changes in microbial communities. In the current study, we conducted high-throughput sequencing of the 16S rRNA gene to analyze the fecal samples from captive infant, young adult, middle-aged, and elderly crab-eating macaques (*Macaca fascicularis*). Our results revealed compositional, functional and network topology changes of gut microbiota associated with its maturation and development. Moreover, our findings identified core age-associated microbes composed of not only commensals but also suspicious pathogens, implicating their importance in the host’s development. We also provided novel evidence supporting a substantial role of driver microbes responsible for age-dependent changes in the gut microbiota network, which were further linked to altered functions of the microbial community. Such findings, taken together, could provide a baseline for better understanding of gut microbiota changes associated with the host’s development and aging in health and diseases.

## Results

### Age-dependent changes of microbiota diversity in healthy captive crab-eating macaques

The metadata of 16s rRNA gene sequencing of fecal DNA was summarized in **Table S1.** Rarefaction analysis of observed operational taxonomic units (OTUs) indicated that the sequencing efficiently captured the potential total OTUs in the fecal samples **(Fig. S1).** The top five phyla observed in the fecal samples of crab-eating macaques were Firmicutes (44.5%-61.1%), Bacteroidetes (26.4%-39.8%), Epsilonbacteraeota (2.3%-8.0%), Proteobacteria (1.9%-3.8%), and Spirochaetes (1.0%-2.7%) **(Fig. 1a)**, with Firmicutes and Bacteroidetes as the two dominant phyla. Furthermore, compared to infants, the Firmicutes to Bacteroidetes (F/B) ratio was found significantly increased in adults (all *P* < 0.05), especially in the middle-aged and elderly. (**Fig. 1b**). The F/B ratio was the lowest in infants (median = 1.09), and increased in young adults (median = 1.28). The highest B/F ratio was observed in the middle-aged (median = 2.74), which slightly decreased in the elderly (median = 2.06) with no significant difference.

**Figure 1.**
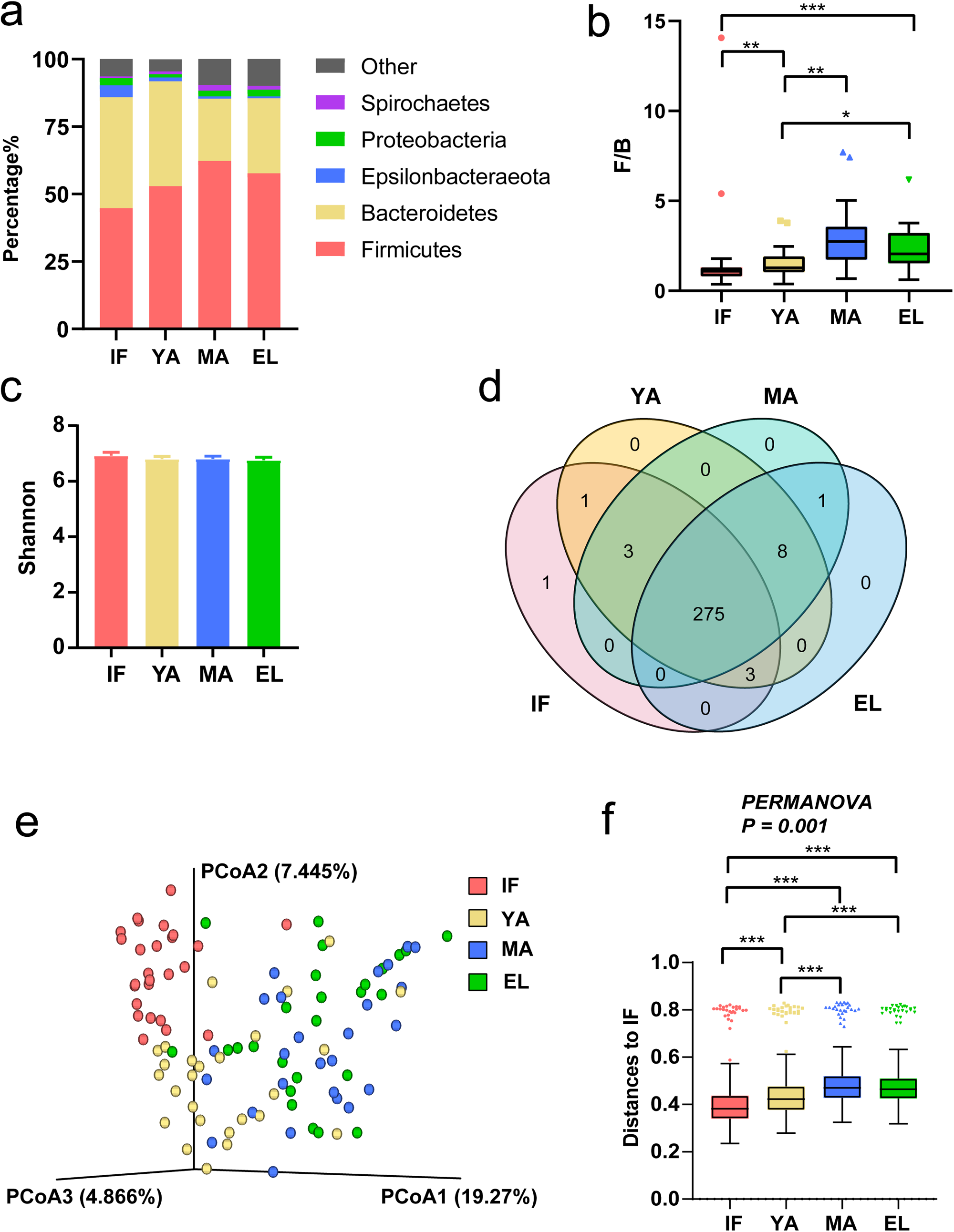
Firmicutes to Bacteroidetes ratio and beta diversity in gut microbiota in different age groups of captive crab-eating macaques. (**a**) Composition of gut microbiota at the phylum level in the age four groups. (**b**) The relative proportion of Firmicutes to Bacteroidetes (F/B) ratio. (**c**) PCoA plot based on the Bray-Curtis distance matrix of all fecal samples. (**d**) Venn plot illustrating overlap of gut microbial genera among age groups. Genera detected in more than 6 fecal samples are included. (**e**) PCoA analysis of all fecal samples based on taxonomic profiles. (**f**) Unweighted Unifrac distance of gut microbiota between the three adult groups and the infant group. Pairwise *P-*values are calculated using nonparametric Kruskal-Wallis test with Tukey post-hoc test. IF, infants; YA, young adults; MA, the middle-aged; EL, the elderly. *: *P*<0.05; **: *P* < 0.01; ***: *P* < 0.001.

Comparison of metrics including the Shannon (**Fig. 1c**) index, Pielou’s evenness, observed OTUs, phylogenetic diversity and Simpson index **(Fig. S2)**, showed no significant change in alpha diversity among the age groups. In line with alpha diversity, the Venn diagram in **Fig. 1d** showed that 275 (94.18%) genera detected in more than six fecal samples were shared across different ages. As for beta diversity, principle coordination analysis (PCoA) based on the Bray-Curtis distance matrix showed that, the infant samples mainly clustered separately from the adult groups **(Fig. 1e)**. The two older adult groups clustered together. The young adult samples fell in-between. Furthermore, permutational multivariate analysis of variance (PERMANOVA) results based on unweighted UniFrac distance indicated significant difference among the four age groups **(Fig. 1f)**. The intergroup unweighted UniFrac distance between adults and infants showed a trend similar to the F/B ratio (median = 0.42, 0.47 and 0.46 in young, middle-aged and elderly adults respectively), compared to the intragroup distance in infants (median = 0.38). These results thus pointed to remarkable microbial community changes associated with age.

### The top abundant gut microbial genera in the four age groups

We then focused on the most abundant genera. Our results showed a trend of age-dependent changes in top abundant genera, similar to that of the beta diversity. The heatmap in **Fig. 2a** showed the top 20 abundant genera from each of the age groups, which were mainly commensals (**Fig. 2b)**. Half of these genera were shared by all age groups (**Fig. 2c**), including four genera from family Ruminococcaceae (*Ruminococcus 1, Ruminococcaceae UCG-005, Ruminococcaceae UCG-014*, and *Subdoligranulum*), three genera from family Prevotellaceae (*Prevotella* 9, *Prevotella* 2, and *Prevotellaceae UCG-003*), *Lactobacillus, Blautia*, and *Dialister*.

**Figure 2.**
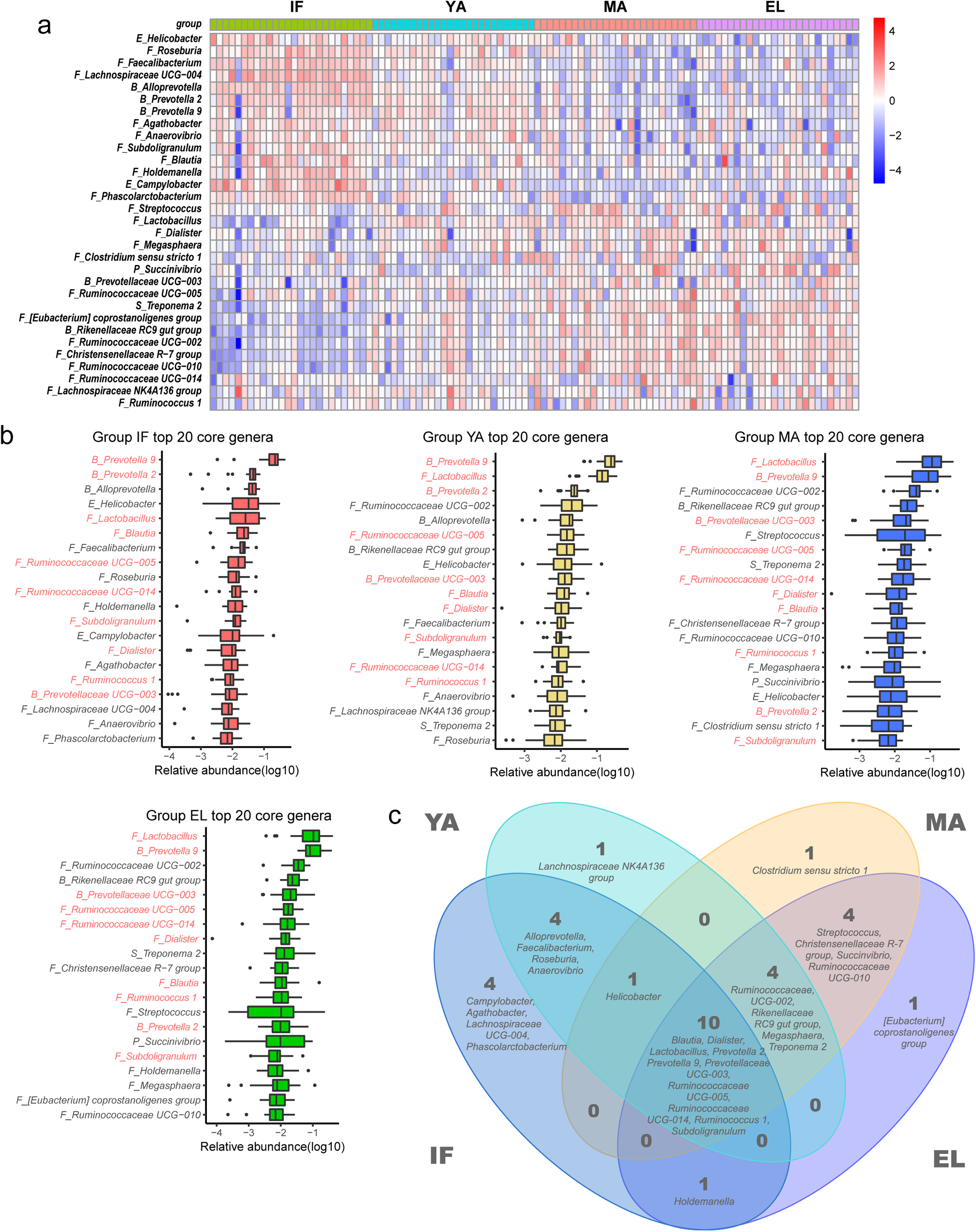
The most abundant genera of gut microbiota in different age groups. (**a**) Heatmap showing the most abundant genera in gut microbiota of the four age groups. (**b**) Box plots showing ranking of top 20 abundant genera in infants, young adults, the middle-aged and elderly. (**c**) Venn plot illustrating overlap of top 20 abundant genera among age groups. Single letters in front of genus names indicate the phylum which the genera belong to: B, *Bacteroidetes*; F, *Firmicutes*; E, *Epsilonbacteraeota*; P, Proteobacteria; S, *Spirochaetes*. IF, infants; YA, young adults; MA, the middle-aged; EL, the elderly.

We also looked into *Bacteroides*, which had been reported to be abundant in gut microbiota of humans living in developed countries [22]. However, the genus show a low mean abundance less than 0.1% in our captive macaques (data not shown).

### Correlation between differentially abundant gut microbes and age

To further characterize age-associated gut microbes, we then identified OTUs with different abundance among age groups using STAMP (**Fig. S3 and S4**). The alluvial plots in **Fig. 3a, 3b, 3c, 3d** and **3e** illustrated clear age-dependent shifts of these taxa at different phylogenetic levels. We further explored their correlation with age using Spearman correlation. At the phylum level **(Fig. 3e and S4)**, *Epsilonbacteraeota, Deferribacteres, Fusobacteria, Bacteroidetes, Patescibacteria*, and *Cyanobacteria* were negatively associated with age, while *Actinobacteria, Kiritimatiellaeota, Lentisphaerae, Firmicutes, WPS-2, Spirochaetes, Planctomycetes, Euryarchaeota*, and *Tenericutes* were negatively associated with age. At the genus level, in total 115 genera were significantly associated with age, with 29 and 18 from family *Lachnospiraceae* and *Ruminococcaceae* respectively (**Fig. S6**). The top 40 genera with the strongest correlations with age were shown in **Fig. 3g**. Among these microbes, 23 genera were negatively associated with age, most of which were potential commensals. These microbes includes night genera from family *Lachnospiraceae* (*Lachnospiraceae UCG-001, Lachnospiraceae UCG-003, Lachnospiraceae UCG-004, Lachnospiraceae UCG-008, [Eubacterium] ventriosum group, Fusicatenibacter, GCA-900066575, [Ruminococcus] torques group*, and *Roseburia*), two genera from family *Prevotellaceae* (*Alloprevotella* and *Prevotella 2*), two genera from family *Ruminococcaceae* (*Faecalibacterium*, and *Fournierella*), *Actinobacillus, Campylobacter, Helicobacter, Mucispirillum, Veillonella, Cetobacterium, Brachyspira*, and *Gemella*. These top age-associated genera also included seventeen genera positively associated with age, including six from the *Ruminococcaceae* family (*Ruminococcaceae UCG-002, Ruminococcaceae UCG-010, Ruminococcaceae UCG-013, Ruminococcaceae NK4A214, CAG-352*, and *[Candidatus] Soleaferrea group*), *Treponema 2, Methanobrevibacter*, the *Rikenellaceae RC9 gut* group, *Christensenellaceae R-7* group, *[Eubacterium] coprostanoligenes group, Lachnospiraceae UCG-007, Libanicoccus, Oscillibacter, Mogibacterium*, and *Stenotrophomonas*.

**Figure 3.**
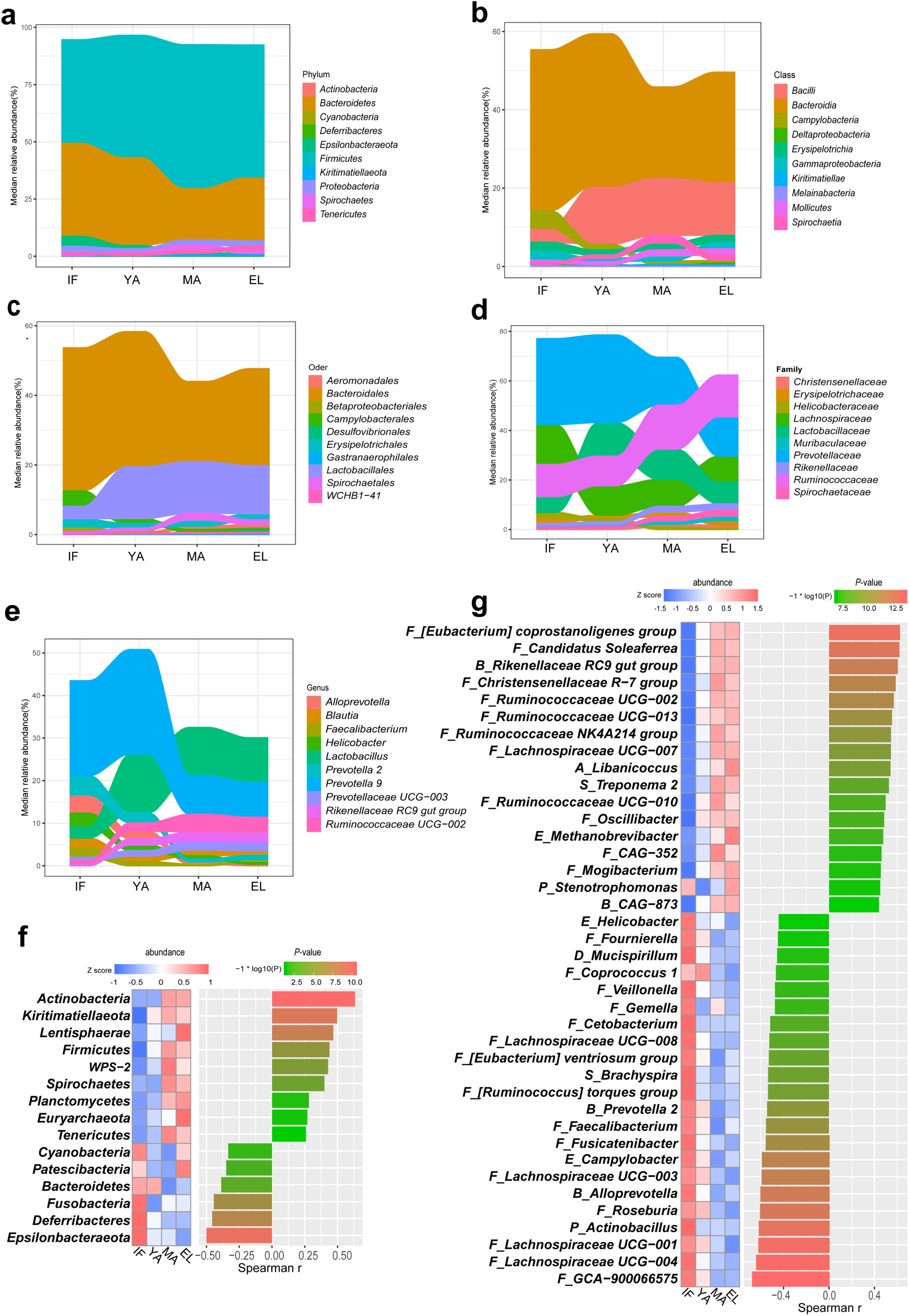
Correlation between differentially abundant gut microbes and age. Alluvial plots illustrating age-dependent phylogenetic shifts of the top 10 differentially abundant taxa at the phylum (**a**), class (**b**), order (**c**), family (**d**) and genus levels (**e**). Differentially abundant taxa are ranked by their median of abundance. Heatmaps showing significant age correlations for differentially abundant phyla (**f**) and genera (**g**) with FDR <0.05. *P-*values are derived from Spearman correlation test. For genera, only the top 40 genera ranked by |r| are shown.

In addition, we also found significantly correlation of with age in lactic acid bacteria known as probiotics in humans (**Fig. S6**). *Bifidobacterium*, which is important in breastfeeding, and *Lactobacillus* that contains the largest number of widely used probiotics, both increased with age (r = 0.34, *P* = 4.2 × 10^−4^ and r = 0.29, *P* = 0.0025 respectively).

### Differential taxa of gut microbiota enriched in the four age groups

We then utilized LEfSe to identify differential taxa most enriched in each of the four age groups. At the phylum level, *Epsilonbacteraeota* and *Cyanobacteria* were enriched in infants, *Firmicutes, Actinobacteria*, and *Kiritimatiellaeota* were enriched in the middle-aged, whereas *Proteobacteria* and *Euryarchaeota* were enriched in the elderly (**Fig 4a**). No phylum was enriched in young adults.

**Figure 4.**
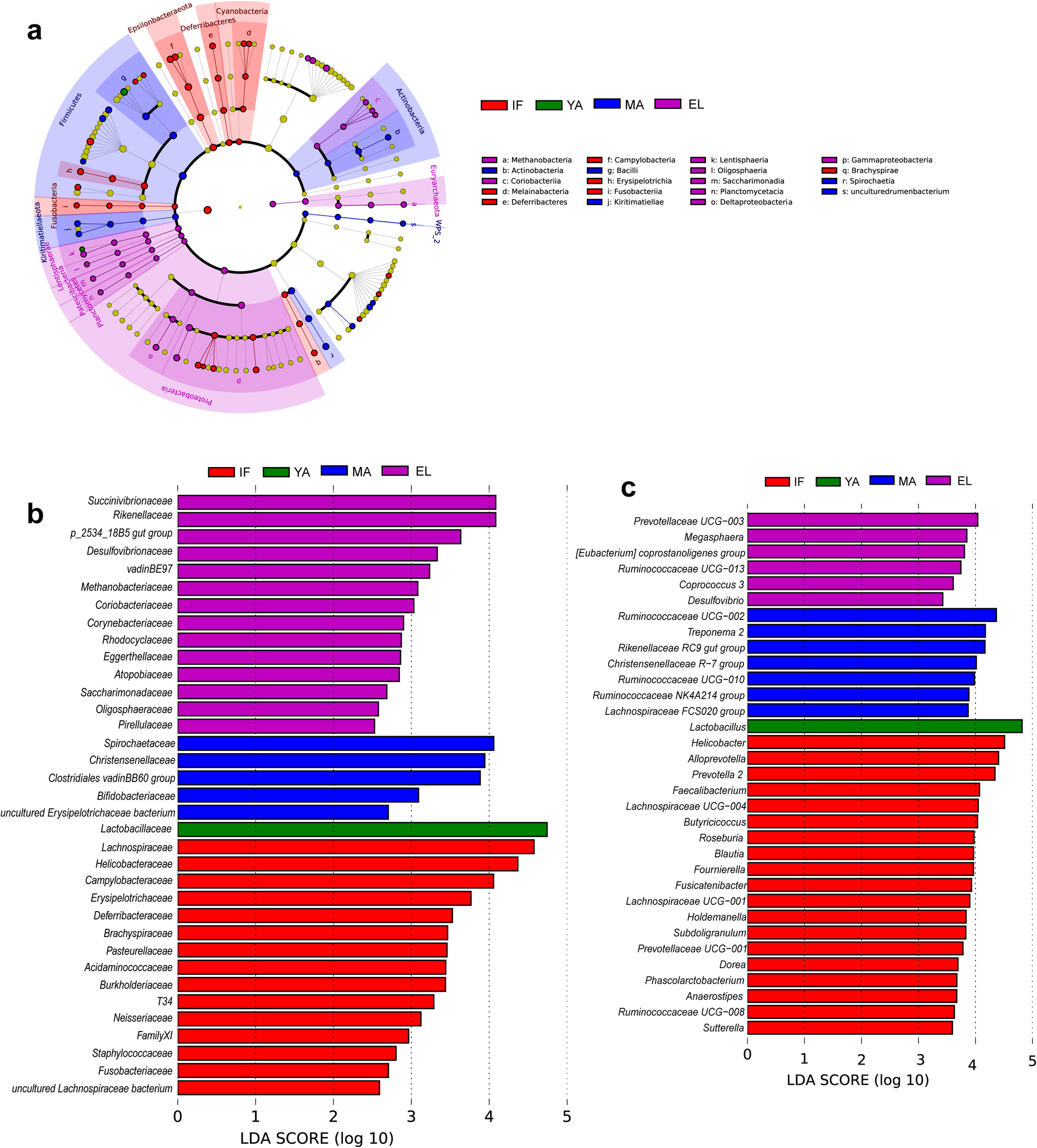
Differentially abundant taxa enriched in the four age groups from LEfSe analysis. (**a**) Phylogenetic cladogram showing differentially abundant taxa from kingdom to family levels. Microbial classes are indicated with letters. Bar charts showing differentially abundant taxa in the family (**b**) and genus levels (**c**) with average abundance > 0.1%.

The largest numbers of enriched families and enriched genera were consistently found in infant macaques (**Fig. 4b**). The family most enriched in infants was *Lachnospiraceae*, and seven of the seventeen infant-enriched genera were from the family, including *Anaerostipes, Blautia, Dorea, Fusicatenibacter, Lachnospiraceae UCG-001, Lachnospiraceae UCG-004*, and *Roseburia. Helicobacter* was the most enriched genus in infants with family *Helicobacteraceae* also enriched in the same group (**Fig. 4c**). Other infant-enriched genera were mainly from family *Prevotellaceae* (*Alloprevotella, Prevotella 2*, and *Prevotellaceae UCG-001*) and family *Ruminococcaceae* (*Butyricicoccus, Faecalibacterium, Fournierella, Ruminococcaceae UCG-008*, and *Subdoligranulum*). Other infant-enriched genera included *Holdemanella* from family *Erysipelotrichaceae, Phascolarctobacterium* from family *Acidaminococcaceae*, and *Sutterella* from *Burkholderiaceae.* In particular, family *Lactobacillaceae* and genus *Lactobacillus*, were uniquely enriched in young adults. It was noticed that *Bifidobacteriaceae*, another group of important lactic acid-producing bacteria were enriched in the middle-aged. Family S*pirochaetaceae* was the most enriched in the middle-aged. Seven genera were enriched in the middle-aged, including three genera from family *Ruminococcaceae* (*Ruminococcaceae NK4A214 group, Ruminococcaceae UCG-002*, and *Ruminococcaceae UCG-010*)), *Treponema 2* from family S*pirochaetaceae, Rikenellaceae RC9 gut group, Christensenellaceae R-7 group*, and *Lachnospiraceae FCS020 group*. In the elderly the most enriched family was *Succinivibrionaceae*. Six genera were enriched in the elderly including *Prevotellaceae UCG_003, Ruminococcaceae UCG-013, Megasphaera* from family *Veillonellaceae, Coprococcus 3* from family *Lachnospiraceae*, and *Desulfovibrio* from *Desulfovibrionaceae*.

### Age-dependent gut microbiota networks and key driver genera

We then further used the Sparse Compositional Correlation (SparCC) analysis to explore the interaction among gut microbes in the four age groups (**Fig. 5**). All genera with relative abundance ≥ 0.1% were included in the networks. Surprisingly, although not preferentially selected, the age-associated genera were found to be the major components of these networks. The gut microbiota network in infants had the lowest connectivity of interactive in infants, as indicated by small Maximal Clique Centrality (MCC) scores (total MCC score = 56) **(Fig. 5a and 6a)**. The network developed into a more mature stage in young adults (total MCC score = 274) **(Fig. 5b and 6a)**, and had the highest connectivity in the middle-aged (total MCC score = 3688) **(Fig. 5c and 6a)**. Unexpectedly, although similar gut microbiota diversities were found between the elderly and middle-aged, the network connectivity dramatically decreased in the elderly (total MCC score = 83) **(Fig. 5d and 6a)**.

**Figure 5.**
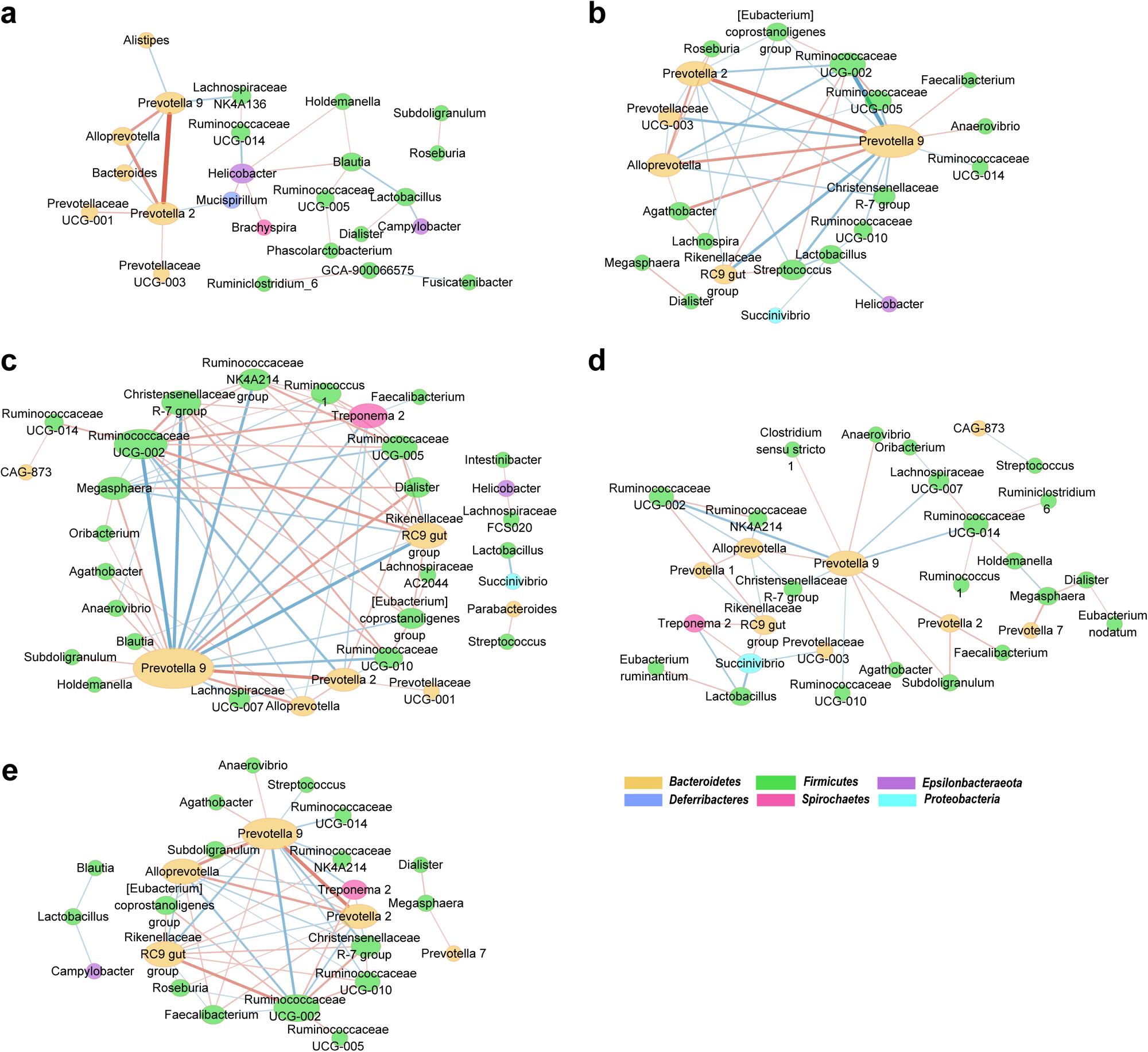
The interactive networks of gut microbiota. Microbial interactive networks in infants (**a**), young adults (**b**), the middle-aged (**c**), the elderly (EL, **d**) and all samples (**e**) are constructed from SparCC results, and visualized using Cytoscape. Genera with average abundance > 0.1%, correlation |r| > 0.2 and *P* < 0.05 are included in the networks. Node colors denote the phylum of the genera. Node sizes represent weighted node connectivity. Edge colors and thickness represent correlation r. IF, infants; YA, young adults; MA, the middle-aged; EL, the elderly.

**Figure 6.**
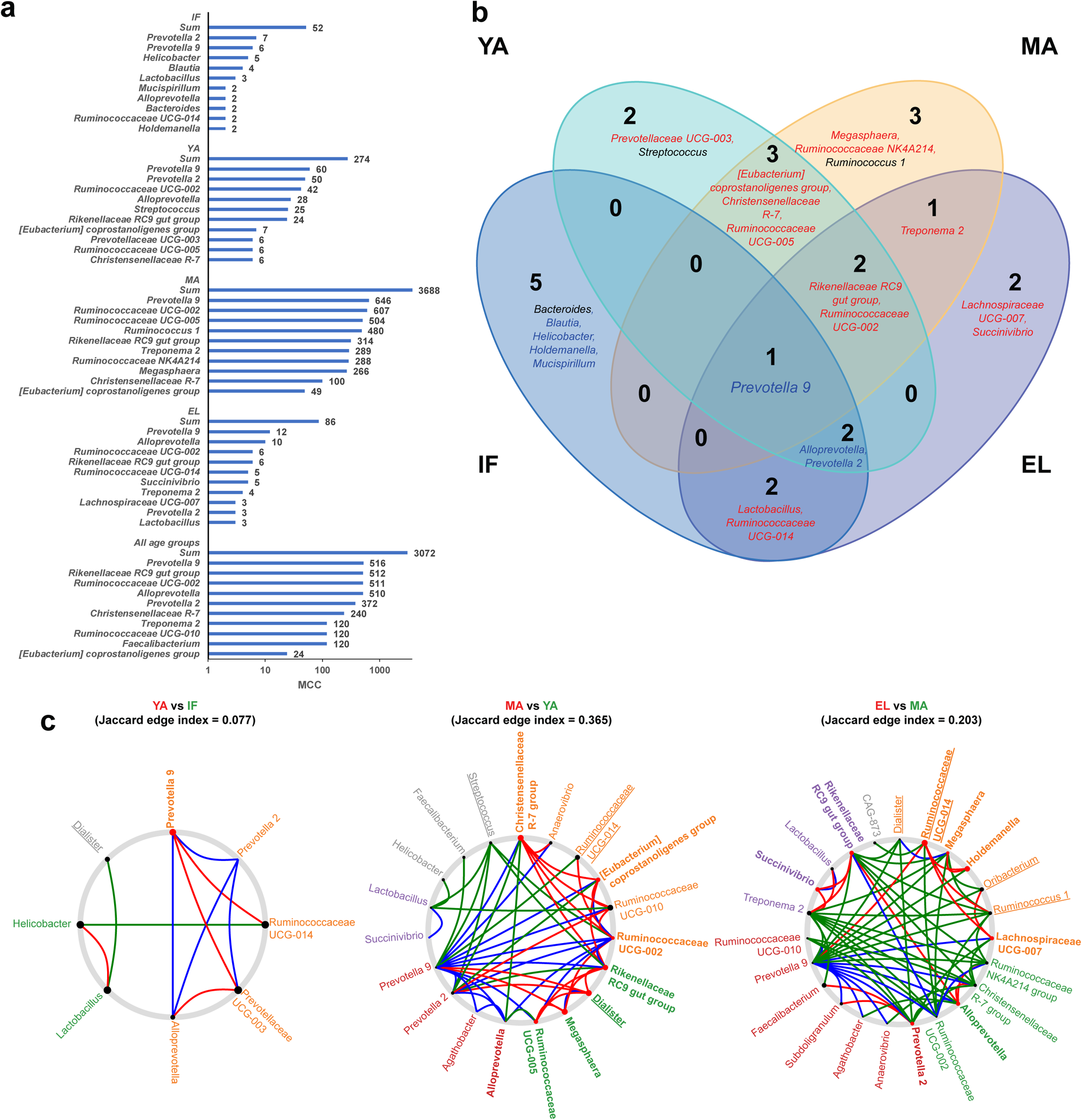
Topological analysis identifies hub and driver genera in the microbiota SparCC networks. (a) MCC scores from the whole network and top 10 hub genera in the SparCC networks. (b) Venn plot showing the overlap of hub genera in the four ages groups. Genera are colored blue if negatively associated with age, and red if positively associated with age. (c) NetShift common sub-networks based on the SparCC networks with highlighted driver genera. Node sizes are in proportion to their NESH scores, and potential drivers are highlighted red. Edges present only in case are colored red, green only in control, and blue in both. Node names without underlines denote age-associated genera.

We then utilized cytoHubba to analyze hub genera, which were supposed to identified by ranking their centralities MCC and EcCentricity (EPC) scores. Among the hub genera shown in **Fig. 6a**, *Prevotella 9* was the only one shared by all four age groups as well as the network constructed using all samples (**Fig. 6a and 6b**). The inter-genera interactions mediated by *Prevotella 9* could be of potential importance. The strongest positive interactions in the microbial communities were found in *Prevotella 2* and *Alloprevotella* with *Prevotella 9* in infants. In addition to *Prevotella 9, Helicobacter* and *Prevotella 2* were another two important hub genera in infants. The role of such interactions mediated by these genera, in particular *Prevotella 9*, gradually diminished with age, and were in part replaced by interactions mediated by hub genera negatively associated with age, such as *Ruminococcaceae UCG-002* and *Rikenellaceae RC9 gut group*.

Moreover, we used NetShift analysis to detect rewiring between microbiota networks, and identified key driver microbes responsible for the changes (**Fig. 6c and Table. S3**). *Prevotella 9* was found to be the only driver genus responsible for the microbial changes between infants and young adults. Novel interactions with *Prevotella 9* were established in the gut microbiota of young adults compared to that of infants. As for adults, multiple potential drivers were identified. Among these drivers, *Rikenellaceae RC9 gut group* and *Megasphaera* are the two key driver genera that contribute to the long-term development of gut microbiota in adults. Another five genera including *Dialister, Christensenellaceae R-7 group, [Eubacterium] coprostanoligenes group, Ruminococcaceae UCG-005* and *Ruminococcaceae UCG-002 group* are involved in the change of gut microbiota between young adults and the middle-aged. Another five genera including *Ruminococcaceae UCG-014, Holdemanella, Succinivibrio, Alloprevotella, Lachnospiraceae UCG-007*, and *Prevotella 2* are involved in the change of gut microbiota between the middle-aged and the elderly.

### Age-associated microbial phenotypes and functions and their correlations with gut microbiota

To understand the potential function impact of age-dependent taxonomic changes in gut microbiota, the microbial phenotypes were predicted using BugBase and compared among age groups. Anaerobic and Gram-positive phenotypes was significantly up-regulated, whereas facultative anaerobic and Gram-negative phenotypes were down-regulated in the middle-aged and elderly groups compared to infants (all *P* < 0.01) (**Fig. 7a**). In line with these findings, Spearman correlation analysis showed that, the anaerobic and Gram-negative phenotypes significantly decreased (*r* = -0.37, *P*=1.2× 10^−4^ and *r* = -0.34, *P* = 4.3× 10^−4^ respectively) with age, whereas the facultative anaerobic and Gram-positive phenotypes significantly increased with age (*r* = 0.42, *P* = *P* = 8.7 × 10^−6^ and *r* = 0.34, *P* = 4.3 × 10^−4^ respectively) (**Fig S6)**.

**Figure 7.**
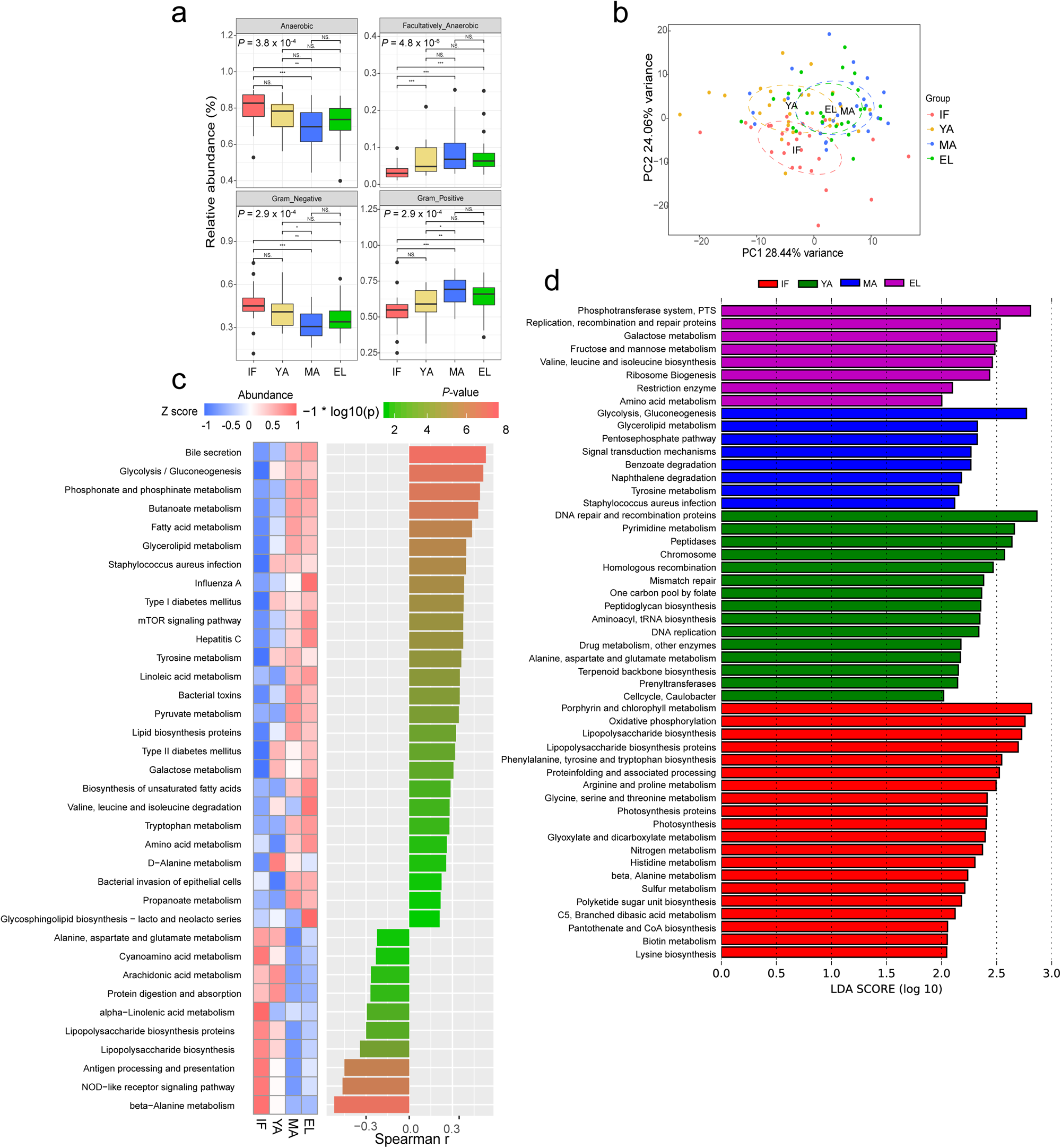
Age-associated gut microbial phenotypes and functional profiles. (a) Comparison of gut microbial phenotypes predicted by BugBase among the four age groups. *P*-values for group comparisons are derived from nonparametric Kruskal-Wallis test with Tukey post-hoc test. (b) PCA plot based on microbial function profiles predicted by PICRUSt. (c) Heatmap illustrating median abundance and age correlation of gut microbial functions related to metabolism of carbohydrates, lipids and proteins as well as host immune response. *P-*values are derived from Spearman correlation test. KEGG pathways with FDR <0.05 are shown. (**d**) LEfSe results of gut microbial functions enriched in each of the four age groups. *: *P*<0.05; **: *P* < 0.01; ***: *P* < 0.001.

We also determined age-dependent changes in gut microbial function using the software Phylogenetic Investigation of Communities by Reconstruction of Unobserved States (PICRUSt), and identified 152 Kyoto Encyclopedia of Genes and Genomes (KEGG) modules to be significantly associated with age (**Table. S2**). The principle component analysis (PCA) plot derived from the abundance of KEGG modules revealed remarkable differences in microbial functions among age groups, showing a similar pattern with beta diversity (**Fig. 7b**). We observed significant correlation between these microbial functions and age. As shown in the heatmap in **Fig. 7c**, metabolic pathways that were the most positively associated with age were mainly involved in biosynthesis and metabolism of lipids, carbohydrates and amino acids. And metabolic pathways that were the most negatively associated with age were mainly involved in host immune response and biosynthesis of the immunomodulating metabolite lipopolysaccharides (LPS), which are endotoxin derived from the outer membrane of Gram-negative bacteria. LEfSe analysis further showed that the pathways related to Porphyrin and chlorophyll metabolism, oxidative phosphorylation and biosynthesis of LPS were upregulated in infants (**Fig. 7d**). In contrast, biosynthesis of peptidoglycan, another important immunomodulating metabolite mainly derived from Gram-positive bacteria, was increased in young adults. Metabolism of carbohydrates was most upregulated in the middle-aged and elderly. Noteworthy, strong correlations were found between these age-associated microbial functions and gut microbes, in particular the hub genera and drivers (**Fig. S8**), with the largest number of positive correlations found in *Prevotella 9*.

## Discussion

By using the NHP model of captive crab-eating macaques, we revealed remarkable lifelong age-dependent changes in gut microbial composition and functions. Moreover, our study identified hub and driver microbes that holds a potential significance in the age-dependent microbial interplay. Given the similarities between the captive crab-eating and humans, these findings could provide better understanding of age-dependent changes in the human gut microbiota.

The gut microbiota of captive macaques in this study showed similarities to that of humans, especially those in developing countries [12, 21, 23, 24]. In line with human and other NHPs, the gut microbiota of our captive crab-eating macaques was dominated by Firmicutes and Bacteroidetes across all ages (**Fig. 1a**) [1, 25]. Most of the common genera with high abundance across all ages, are potentially commensals from the Ruminococcaceae and Prevotellaceae families such as *Prevotella 9* (**Fig. 2b**). In contrast, *Bacteroides* had very low abundance. Gut microbial communities of individuals from developing countries has been reported to be dominated by *Prevotella*[24], while those from developed countries was highly abundant in *Bacteroides*[26]. Plant-based diets with low fat could be involved in the higher similarities between the gut microbiota of and captive macaques and humans living in developing countries [22].

The lack of significant change in alpha diversity might indicate the important of microbiota studies in captive NHPs (**Fig. 1**). Yatsunenko *et al.* reported that observed OTUs increased with age in gut microbiota of all three populations [12]. In a recent gut microbiota study of non-captive rhesus macaques Chen *et al.* reported that male adults had significant higher Shannon index than male juvenile [27]. However, under a well-controlled environment provided by captivity, alpha diversity changes are probably smoothed out. By age of 1-2 years old, infant gut microbiota had gained more than 94% of OTUs observed in adults (**Fig 2a**). Age-related factors, such as diets and life styles, rather than age itself, might actually contribute to the age-associated increase of alpha diversity in human populations.

Nevertheless, the remarkable age-dependent changes including the F/B ratio and beta diversity as well as network topology emphasized actual effects of age on the gut microbiota in captive macaques (**Fig. 1b**). The F/B ratio is considered as an indicator of maturation and development of gut microbiota [28], and has been reported to be involved in health-related conditions or diseases such as obesity [29]. In the current study the F/B ratio increased in adult macaques, and decreased in elderly macaques (**Fig. 1b**), resembling observation in humans [28, 30]. It could be due to increased Firmicutes and decreased Bacteroidetes with age (**Fig. 3f**). Interestingly, although middle-aged and elderly macaques had similar beta diversity, evident reduction of connectivity in elderly macaques, indicating a decline of microbial interactions. Such findings suggest that, network connectivity could be more sensitive than the F/B ratio and biological diversity to detect age-dependent changes in the gut microbiota.

Moreover, the age-associated microbes identified in captive macaques could be involved in the host’s development and aging in good health (**Fig. 3 and 4**). These microbes could play distinct roles dependent of their direction of age-correlation. A large proportion of these age-associated genera decreased with age, including those enriched in infants. The composition and activities in the infant gut microbiota has been engaged in the host’s early development and a variety of diseases, such as allergy and autisms [5, 31, 32]. These genera negatively associated with age in fact consisted of at least two distinct groups. First, these genera contained potential commensals, which were active players in the early development of gut microbiota (**Fig. 4b, S4, and S6**). The interplay between these commensals and the host intestinal barriers are important to the postnatal development of host metabolism, immunity and mucosal barrier [33-35]. Commensals could benefit the host by producing metabolites such as short chain fatty acids [36]. A number of the age-associated commensals in the current study are butyrate-producing bacteria in the host colon, including *Faecalibacterium, Roseburia, Anaerostipes*, and *Butyricicoccus* [37]. These commensals include anti-inflammatory bacteria, and outcompete pathogens to protect the host, and abnormal alteration of them has been reported in various human diseases [38-42]. For example, *Faecalibacterium prausnitzii* is one of the most abundant anti-inflammatory commensal bacteria in the colon, and was reduced in Crohn disease patients [40].

Second, these bacteria negative associated with age also contained a number of suspicious pathogens, especially enteropathogens (**Fig. 4b, S4, and S6**). *Campylobacter* and *Actinobacillus* are causes of infectious diseases in humans, *Campylobacteriosis* and *Actinobacillosis* [43]. Species from the genus *Brachyspira* are known pathogens causing diarrhea in animals and human [44]. Bacteria from the *Gemella* genus are involved in endocarditis [45]. *Anaerobiospirillum succiniciproducens* from the genus *Anaerobiospirillum* has been found to be associated with has diarrhea and bacteremia [45]. It was noted that *Helicobacter*, a group of Gram-negative bacteria, was identified as a hub genus with high abundance in infant gut microbiota, but its role remained largely unclear. *Helicobacter macacae* from the genus have been reported to be frequently detected in rhesus monkeys without a diarrheal history [46]. Rhoades et al. report that 8-month infant remained asymptomatic for diarrhea were enriched for the species [9]. In line with these findings, biosynthesis of LPS was also upregulated in our infant macaques (**Fig.7**), further supporting a potential role of these age-associated microbes in modulation of the host’s immunity. It should be taken into account that all macaques in the current study were in good health. Therefore, the gradual decrease of these suspicious pathogens with age might associated with the maturation of gut mucosal barrier. In addition, recent studies have reported possible effects of pathogens protecting the host against allergic sensitization [47, 48]. In our captive macaques the suspicious pathogens with their abundance under control might allow “good” exposure for the proper training of the host’s immune system.

While the roles of the microbes positively associated with age remained largely unclear, they could be related to the host’s healthy aging (**Fig. 4b, S4, and S6**). A subset of these microbes has been implicated to be involved in metabolism of nutrients including lipids and carbohydrates, which are in line with the predicted gut microbial functions upregulated with age in our macaques. Importantly, the genus *Lactobacillus*, highly abundant in our adult macaques (**Fig. 2**), are widely used probiotics with potential effects on lipid metabolism [49]. We also notice that *Bifidobacterium*, the key probiotics for the metabolism of oligosaccharides in breast milk [50], also increased with age. The increase of these lactic acid-producing probiotics might indicate a potential role of these bacteria in healthy aging. In addition, *Eubacterium coprostanoligenes* had been identified as a cholesterol-reducing anaerobe [51]. genera *Christensenellaceae R-7 group, Ruminococcaceae* (*UCG-002*, and *UCG-010*), and *Lachnospiraceae FCS020 group* were linked to circulating lipid-related metabolites in a recent population-based study[52]. *Candidatus soleaferrea* was increased in a randomized controlled trial of hypocaloric diet with Hass avocado [53]. In line with these findings, changes of microbial functions related to metabolisms of lipids and carbohydrates increased with age (**Fig. 7b**). In addition, these microbes positively associated with age are also involved in diseases. *Treponema 2, Rikenellaceae RC9 gut group, Prevotellaceae UCG-003* were increased in rats with isoproterenol-induced acute myocardial ischemia[54], whereas in a meta-analysis *Christensenellaceae R-7 group* was found to be reduced in patients affected by intestinal diseases [55]. Intriguingly, although the reported role of archaea in the host’s health remain unclear, our results showed that, the archaeal family *Methanobacteriaceae* was differentially enriched in elderly macaques, and genus *Methanobrevibacter* increased with age in the macaque gut microbiota. Such findings thus indicate a positive association of these methanogens with host aging.

This study further highlights the pivotal role of driver microbes in age-dependent changes of the gut microbiota (**Fig. 5 and 6**). Genus *Prevotella 9*, with a high abundance in our captive macaques, was identified as the most important hub mediating large proportion of microbial interactions in gut microbiotas across all ages. And it acted as the key driver responsible for the gut microbiota maturation from infants to young adults. The exact biological significance of *Prevotella 9* in the context of integrative bacterial community and microbiota development has yet to be further elucidated. A recent reanalysis of existing gut metagenomes from NHPs and humans reported that *Prevotella* were prevalent in primate gut microbiota of different host species [20]. In line with such finding, the *Prevotella 9* genus was highly abundant across all ages with gradual age-dependent decrease in our captive macaques. The high abundance of the genus in primates could be strongly associated with plant-based, low-fat diets [22]. In addition, the high abundance of *Prevotella* in humans and NHPs might also have possible implications for host-microbiota coevolution [56]. Although *Prevotella* 9 remained abundant in adult macaques, its level decreased with age, and possibly freed up space for other microbes that were necessary for further microbiota development, such as *Rikenellaceae RC9 gut group* and *Megasphaera*. Such shift of driver microbes could in turn impact the changes of gut microbiota phenotypes and functions.

## Conclusions

In summary, by using captive crab-eating macaques to control confounding factors, the current study demonstrates evident age-dependent structural and functional changes in the healthy gut microbiota during the host’s development and aging. Our key findings of age-associated microbes, composed of both commensals and suspicious pathogens, indicate the potential importance of appropriate bacterial exposure for the early development of the host. Moreover, the hub and driver microbes identified by network topology analysis probably play a pivotal role as core microbes in the microbial communities, and are responsible for the maturation and development of the primate gut microbiota. By characterizing the age-dependent changes in the gut microbiota during the host’s development and healthy aging, the current study also provides a baseline for comparison and understanding of the role of the primate gut microbiota in health and disease.

## Materials and methods

### Animals in the study

A total of 104 male crab-eating macaques from Guangdong Xiangguan Biotechnology Co. Ltd. (Guangzhou, China) were included in the current study. All of the animals were confirmed to be in good health by records and veterinary examination prior to the study. These animals were composed of four different age-groups (N=26 for each group), including infant (1-2 years old), young adult (4-6 years old), middle-aged group (8-10 years old), and an elderly macaques (≥13 years old). Post-weaning infant macaques were selected to reduce possible effects of breastfeeding. All animals were kept in a well-controlled environment with moderate room temperature (16-28 °C) and relative humidity of 40%-70%, as well as a 12/12-hour light-dark cycle. The study complied with protocols approved by the Animal Ethics Committees of Guangdong Institute of Applied Biological Resources, and were in compliance with the Guide for the Care and Use of Laboratory Animals [57].

### Stool sample collection and DNA extraction

Rectal swab samples were freshly collected from each monkey, and stored at -80 °C immediately until DNA extraction. Microbial DNA was extracted using TIANamp Stool DNA kit (Cat.#DP328, Tiangen, China) according to the manufacturer’s instructions, and its concentration and quality were assessed using a Nanodrop One Microvolume UV Spectrophotometer (Thermofisher, U.S.).

### 16S rRNA gene sequencing

The hypervariable V4 regions of bacterial/archaeal 16s rRNA genes were amplified using polymerase chain reaction and V4-specific primers 515F (5′-GTGCCAGCMGCCGCGGTAA-3′) and 806R (5′-GGACTACHVGGGTWTCTAAT-3′). PCR products between 400 and 450 bp were checked using the 2% agarose gel, purified using GeneJET Gel Extraction Kit (Thermo Fisher Scientific, USA), and sequenced on an Ion S5XL sequencer with a single-end 400-bp read length configuration.

### Processing of 16S rRNA gene sequencing data

Bioinformatic analysis of the 16S rRNA gene sequencing data was performed using the QIIME2 (version 2018.6.0) analysis pipeline [58]. Briefly, sequencing data were processed by the dada2 program to filter low-quality and chimeric sequences, and generate unique feature tables equivalent to OTU tables at exact match or 100% sequence similarity. Taxonomy was then assigned to these features using the q2-feature-classifier against the full-length SILVA database (release r132) at 99% similarity cutoff [59]. Analysis of microbiota diversities were conducted in QIIME2: alpha diversity metrics including Pielou’s evenness, phylogenetic diversity, observed OTUs, Shannon and Simpson’s indices, and beta diversity including weighted/unweighted UniFrac distances, and Bray-Curtis dissimilarity. Comparison of beta diversity was performed using the nonparametric method PERMANOVA. Abundance of OUTs were compared among groups by using STAMP [59]. the Linear discriminant analysis (LDA) Effect Size (LEfSe) algorithm was used with a log (LDA) score cutoff of 2 to identify taxa specifically enriched in particular age groups [60]. Phylogenetic cladograms of LEfSe results were visualized using the GraPhlAn tool (https://bitbucket.org/nsegata/graphlan).

### Microbial interactive network construction and analysis

The SparCC (https://bitbucket.org/yonatanf/sparcc) algorithm was used to estimate the correlations among gut microbes [61]. 100 bootstrap replicates were used to calculate the pseudo *P*-values in the SparCC analysis, and correlations with | correlation coefficient (r) | >0.2 and *P* < 0.01 were considered significant. For each OTU with significant SparCC correlation, a weighted node connectivity score was calculated as an indicator of its weight in the network, by summing up its | r | with all of its first neighbors [62]. The constructed gut microbial interactive network was further visualized using Cytoscape version 3.7.0 [63]. The cytoHubba plugin was used to identify hub genera in the networks [64]. Two node ranking methods including a local-based method MCC and a global-based method EPC were used to evaluate importance of genera. In addition, NetShift (https://web.rniapps.net/netshift/) was used to evaluate potential driver microbes using a case-control strategy to compare a pair of networks as described [64, 65]. Neighbor Shift (NESH) Scores were calculated to quantify enriched interaction in the case over the control.

### Prediction of microbial phenotypes and function profiles

The BugBase (https://bugbase.cs.umn.edu/) analysis tool was utilized to predict high-level phenotypes in fecal microbiome samples. PICRUSt version 1.1.4 was used to predict microbial functions from the 16S rRNA gene sequencing data, which were further categorized using the BRITE hierarchy of the KEGG database [66]. PCA based on KEGG module abundance was conducted using STAMP.

### Statistical analysis

Statistical analysis was performed using GraphPad Prism V.7.0a (GraphPad Software, USA) and the R statistical language (version 3.6.0). Abundance of OTUs and KEGG modules among groups were compared using the non-parametric Kruskal-Wallis test, and evaluated for pair-wise inter-group differences with Tukey’s post hoc test if overall significance was found. The Benjamini-Hochberg false discovery rate (FDR) correction was applied for multiple testing. Correlations of OTUs, microbial phenotypes and KEGG functions with age were determined using Spearman’s correlation analysis. Differences in the taxa were analyzed by LEfSe with default settings.

## Ethics approval and consent to participate

The study complied with protocols approved by the Animal Ethics Committees of Guangdong Institute of Applied Biological Resources, and were in compliance with the Guide for the Care and Use of Laboratory Animals.

## Consent for publication

Not applicable.

## Data availability

The raw datasets of 16s rRNA gene amplicon sequencing in the current study are deposited and available in the BioProject (https://www.ncbi.nlm.nih.gov/bioproject) repository under the accession number PRJNA598010.

## Authors’ contributions

Z-YW and J-HR contributed equally to this work. J-HR and J-HC conceived the project and planned the experiments. J-HR, B-HL and M-TT collected the fecal samples. Z-YW, M-TT, G-AZ, Q-CL, L-MW, B-QX performed the experiments. Z-YW, G-AZ, X-YL and J-HC analyzed and interpreted the experiment data. Z-YW, X-YL and J-HC drafted the manuscript. All authors read and approved the final manuscript.

## Acknowledgement

Not applicable.

## References

[1] Turnbaugh PJ, Ley RE, Hamady M, Fraser-Liggett CM, Knight R, Gordon JI. The human microbiome project. Nature 2007;449:804–10.

[2] Gill SR, Pop M, Deboy RT, Eckburg PB, Turnbaugh PJ, Samuel BS, et al. Metagenomic analysis of the human distal gut microbiome. Science 2006;312:1355–9.

[3] Shreiner AB, Kao JY, Young VB. The gut microbiome in health and in disease. Curr Opin Gastroenterol 2015;31:69–75.

[4] Rehman T. Role of the Gut Microbiota in Age-Related Chronic Inflammation. Endocrine, Metabolic & Immune Disorders-Drug Targets 2012;12:361–7.

[5] Buford TW. (Dis)Trust your gut: the gut microbiome in age-related inflammation, health, and disease. Microbiome 2017;5:80.

[6] Arpaia N, Campbell C, Fan X, Dikiy S, van der Veeken J, deRoos P, et al. Metabolites produced by commensal bacteria promote peripheral regulatory T-cell generation. Nature 2013;504:451–5.

[7] O’Toole PW, Jeffery IB. Gut microbiota and aging. Science 2015;350:1214–5.

[8] Milani C, Duranti S, Bottacini F, Casey E, Turroni F, Mahony J, et al. The First Microbial Colonizers of the Human Gut: Composition, Activities, and Health Implications of the Infant Gut Microbiota. Microbiol Mol Biol Rev 2017;81.

[9] Rhoades N, Barr T, Hendrickson S, Prongay K, Haertel A, Gill L, et al. Maturation of the infant rhesus macaque gut microbiome and its role in the development of diarrheal disease. Genome Biology 2019;20:173.

[10] Hill CJ, Lynch DB, Murphy K, Ulaszewska M, Jeffery IB, O’Shea CA, et al. Evolution of gut microbiota composition from birth to 24 weeks in the INFANTMET Cohort. Microbiome 2017;5:4.

[11] Garrido D, Ruiz-Moyano S, Jimenez-Espinoza R, Eom HJ, Block DE, Mills DA. Utilization of galactooligosaccharides by Bifidobacterium longum subsp. infantis isolates. Food Microbiol 2013;33:262–70.

[12] Yatsunenko T, Rey FE, Manary MJ, Trehan I, Dominguez-Bello MG, Contreras M, et al. Human gut microbiome viewed across age and geography. Nature 2012;486:222–7.

[13] Li J, Zhao F, Wang Y, Chen J, Tao J, Tian G, et al. Gut microbiota dysbiosis contributes to the development of hypertension. Microbiome 2017;5:14.

[14] Qin J, Li Y, Cai Z, Li S, Zhu J, Zhang F, et al. A metagenome-wide association study of gut microbiota in type 2 diabetes. Nature 2012;490:55–60.

[15] David LA, Maurice CF, Carmody RN, Gootenberg DB, Button JE, Wolfe BE, et al. Diet rapidly and reproducibly alters the human gut microbiome. Nature 2014;505:559–63.

[16] Perez-Cobas AE, Gosalbes MJ, Friedrichs A, Knecht H, Artacho A, Eismann K, et al. Gut microbiota disturbance during antibiotic therapy: a multi-omic approach. Gut 2013;62:1591–601.

[17] Rinninella E, Raoul P, Cintoni M, Franceschi F, Miggiano GAD, Gasbarrini A, et al. What is the Healthy Gut Microbiota Composition? A Changing Ecosystem across Age, Environment, Diet, and Diseases. Microorganisms 2019;7.

[18] Dominguez-Bello MG, Godoy-Vitorino F, Knight R, Blaser MJ. Role of the microbiome in human development. Gut 2019;68:1108–14.

[19] Walter J, Armet AM, Finlay BB, Shanahan F. Establishing or Exaggerating Causality for the Gut Microbiome: Lessons from Human Microbiota-Associated Rodents. Cell 2020;180:221–32.

[20] Manara S, Asnicar F, Beghini F, Bazzani D, Cumbo F, Zolfo M, et al. Microbial genomes from non-human primate gut metagenomes expand the primate-associated bacterial tree of life with over 1000 novel species. Genome Biol 2019;20:299.

[21] Clayton JB, Vangay P, Huang H, Ward T, Hillmann BM, Al-Ghalith GA, et al. Captivity humanizes the primate microbiome. Proc Natl Acad Sci U S A 2016;113:10376–81.

[22] Ou J, Carbonero F, Zoetendal EG, DeLany JP, Wang M, Newton K, et al. Diet, microbiota, and microbial metabolites in colon cancer risk in rural Africans and African Americans. Am J Clin Nutr 2013;98:111–20.

[23] Yasuda K, Oh K, Ren B, Tickle TL, Franzosa EA, Wachtman LM, et al. Biogeography of the intestinal mucosal and lumenal microbiome in the rhesus macaque. Cell Host Microbe 2015;17:385–91.

[24] Lin A, Bik EM, Costello EK, Dethlefsen L, Haque R, Relman DA, et al. Distinct distal gut microbiome diversity and composition in healthy children from Bangladesh and the United States. PLoS One 2013;8:e53838.

[25] Mahowald MA, Rey FE, Seedorf H, Turnbaugh PJ, Fulton RS, Wollam A, et al. Characterizing a model human gut microbiota composed of members of its two dominant bacterial phyla. Proc Natl Acad Sci U S A 2009;106:5859–64.

[26] Yatsunenko T, Rey FE, Manary MJ, Trehan I, Dominguez-Bello MG, Contreras M, et al. Human gut microbiome viewed across age and geography. Nature 2012;486:222–7.

[27] Chen Z, Yeoh YK, Hui M, Wong PY, Chan MCW, Ip M, et al. Diversity of macaque microbiota compared to the human counterparts. Sci Rep 2018;8:15573.

[28] Mariat D, Firmesse O, Levenez F, Guimaraes V, Sokol H, Dore J, et al. The Firmicutes/Bacteroidetes ratio of the human microbiota changes with age. BMC Microbiol 2009;9:123.

[29] Koliada A, Syzenko G, Moseiko V, Budovska L, Puchkov K, Perederiy V, et al. Association between body mass index and Firmicutes/Bacteroidetes ratio in an adult Ukrainian population. BMC Microbiol 2017;17:120.

[30] Claesson MJ, Cusack S, O’Sullivan O, Greene-Diniz R, de Weerd H, Flannery E, et al. Composition, variability, and temporal stability of the intestinal microbiota of the elderly. Proc Natl Acad Sci U S A 2011;108 Suppl 1:4586–91.

[31] Rodríguez JM, Murphy K, Stanton C, Ross RP, Kober OI, Juge N, et al. The composition of the gut microbiota throughout life, with an emphasis on early life. Microbial Ecology in Health & Disease 2015;26.

[32] Sharon G, Cruz NJ, Kang DW, Gandal MJ, Wang B, Kim YM, et al. Human Gut Microbiota from Autism Spectrum Disorder Promote Behavioral Symptoms in Mice. Cell 2019;177:1600–18 e17.

[33] Tlaskalova-Hogenova H, Stepankova R, Kozakova H, Hudcovic T, Vannucci L, Tuckova L, et al. The role of gut microbiota (commensal bacteria) and the mucosal barrier in the pathogenesis of inflammatory and autoimmune diseases and cancer: contribution of germ-free and gnotobiotic animal models of human diseases. Cell Mol Immunol 2011;8:110–20.

[34] Lopez-Siles M, Duncan SH, Garcia-Gil LJ, Martinez-Medina M. Faecalibacterium prausnitzii: from microbiology to diagnostics and prognostics. ISME J 2017;11:841–52.

[35] Hsiao A, Ahmed AM, Subramanian S, Griffin NW, Drewry LL, Petri WA, Jr., et al. Members of the human gut microbiota involved in recovery from Vibrio cholerae infection. Nature 2014;515:423–6.

[36] Martin R, Miquel S, Ulmer J, Kechaou N, Langella P, Bermudez-Humaran LG. Role of commensal and probiotic bacteria in human health: a focus on inflammatory bowel disease. Microb Cell Fact 2013;12:71.

[37] Riviere A, Selak M, Lantin D, Leroy F, De Vuyst L. Bifidobacteria and Butyrate-Producing Colon Bacteria: Importance and Strategies for Their Stimulation in the Human Gut. Front Microbiol 2016;7:979.

[38] Abt MC, Pamer EG. Commensal bacteria mediated defenses against pathogens. Curr Opin Immunol 2014;29:16–22.

[39] Herp S, Brugiroux S, Garzetti D, Ring D, Jochum LM, Beutler M, et al. Mucispirillum schaedleri Antagonizes Salmonella Virulence to Protect Mice against Colitis. Cell Host & Microbe 2019;25:681-94.e8.

[40] Munukka E, Rintala A, Toivonen R, Nylund M, Yang B, Takanen A, et al. Faecalibacterium prausnitzii treatment improves hepatic health and reduces adipose tissue inflammation in high-fat fed mice. The ISME Journal 2017;11:1667–79.

[41] Jenq RR, Taur Y, Devlin SM, Ponce DM, Goldberg JD, Ahr KF, et al. Intestinal Blautia Is Associated with Reduced Death from Graft-versus-Host Disease. Biol Blood Marrow Transplant 2015;21:1373–83.

[42] Eeckhaut V, Machiels K, Perrier C, Romero C, Maes S, Flahou B, et al. Butyricicoccus pullicaecorum in inflammatory bowel disease. Gut 2013;62:1745–52.

[43] Coker AO, Isokpehi RD, Thomas BN, Amisu KO, Obi CL. Human Campylobacteriosis in Developing Countries1. Emerging Infectious Diseases 2002;8:237–43.

[44] Mirajkar NS, Phillips ND, La T, Hampson DJ, Gebhart CJ. Characterization and Recognition of Brachyspira hampsonii sp. nov., a Novel Intestinal Spirochete That Is Pathogenic to Pigs. Journal of Clinical Microbiology 2016;54:2942–9.

[45] La Scola B, Raoult D. Molecular identification of Gemella species from three patients with endocarditis. J Clin Microbiol 1998;36:866–71.

[46] Marini RP, Muthupalani S, Shen Z, Buckley EM, Alvarado C, Taylor NS, et al. Persistent infection of rhesus monkeys with ‘Helicobacter macacae’ and its isolation from an animal with intestinal adenocarcinoma. J Med Microbiol 2010;59:961–9.

[47] Lynch SV, Wood RA, Boushey H, Bacharier LB, Bloomberg GR, Kattan M, et al. Effects of early-life exposure to allergens and bacteria on recurrent wheeze and atopy in urban children. J Allergy Clin Immunol 2014;134:593–601 e12.

[48] Reynolds LA, Finlay BB. Early life factors that affect allergy development. Nature Reviews Immunology 2017;17:518–28.

[49] Xie N, Cui Y, Yin YN, Zhao X, Yang JW, Wang ZG, et al. Effects of two Lactobacillus strains on lipid metabolism and intestinal microflora in rats fed a high-cholesterol diet. BMC Complement Altern Med 2011;11:53.

[50] Gueimonde M, Laitinen K, Salminen S, Isolauri E. Breast milk: a source of bifidobacteria for infant gut development and maturation? Neonatology 2007;92:64–6.

[51] Freier TA, Beitz DC, Li L, Hartman PA. Characterization of Eubacterium coprostanoligenes sp. nov., a cholesterol-reducing anaerobe. Int J Syst Bacteriol 1994;44:137–42.

[52] Vojinovic D, Radjabzadeh D, Kurilshikov A, Amin N, Wijmenga C, Franke L, et al. Relationship between gut microbiota and circulating metabolites in population-based cohorts. Nat Commun 2019;10:5813.

[53] Henning SM, Yang J, Woo SL, Lee RP, Huang J, Rasmusen A, et al. Hass Avocado Inclusion in a Weight-Loss Diet Supported Weight Loss and Altered Gut Microbiota: A 12-Week Randomized, Parallel-Controlled Trial. Curr Dev Nutr 2019;3:nzz068.

[54] Sun L, Jia H, Li J, Yu M, Yang Y, Tian D, et al. Cecal Gut Microbiota and Metabolites Might Contribute to the Severity of Acute Myocardial Ischemia by Impacting the Intestinal Permeability, Oxidative Stress, and Energy Metabolism. Front Microbiol 2019;10:1745.

[55] Mancabelli L, Milani C, Lugli GA, Turroni F, Cocconi D, van Sinderen D, et al. Identification of universal gut microbial biomarkers of common human intestinal diseases by meta-analysis. FEMS Microbiol Ecol 2017;93.

[56] Ochman H, Worobey M, Kuo CH, Ndjango JB, Peeters M, Hahn BH, et al. Evolutionary relationships of wild hominids recapitulated by gut microbial communities. PLoS Biol 2010;8:e1000546.

[57] National Research Council (U.S.). Committee for the Update of the Guide for the Care and Use of Laboratory Animals., Institute for Laboratory Animal Research (U.S.), National Academies Press (U.S.) (2011), ‘Guide for the care and use of laboratory animals’, National Academies Press,, Washington, D.C., pp. xxv, 220 p.

[58] Bolyen E, Rideout JR, Dillon MR, Bokulich NA, Abnet CC, Al-Ghalith GA, et al. Reproducible, interactive, scalable and extensible microbiome data science using QIIME 2. Nat Biotechnol 2019;37:852–7.

[59] Parks DH, Tyson GW, Hugenholtz P, Beiko RG. STAMP: statistical analysis of taxonomic and functional profiles. Bioinformatics 2014;30:3123–4.

[60] Segata N, Izard J, Waldron L, Gevers D, Miropolsky L, Garrett WS, et al. Metagenomic biomarker discovery and explanation. Genome Biol 2011;12:R60.

[61] Friedman J, Alm EJ. Inferring Correlation Networks from Genomic Survey Data. PLOS Computational Biology 2012;8:11.

[62] Azuaje FJ. Selecting biologically informative genes in co-expression networks with a centrality score. Biology Direct 2014;9:12.

[63] Shannon P, Markiel A, Ozier O, Baliga NS, Wang JT, Ramage D, et al. Cytoscape: a software environment for integrated models of biomolecular interaction networks. Genome Res 2003;13:2498–504.

[64] Chin C-H, Chen S-H, Wu H-H, Ho C-W, Ko M-T, Lin C-Y. cytoHubba: identifying hub objects and sub-networks from complex interactome. BMC Systems Biology 2014;8:S11.

[65] Kuntal BK, Chandrakar P, Sadhu S, Mande SS. ‘NetShift’: a methodology for understanding ‘driver microbes’ from healthy and disease microbiome datasets. The ISME Journal 2019;13:442–54.

[66] Langille MGI, Zaneveld J, Caporaso JG, McDonald D, Knights D, Reyes JA, et al. Predictive functional profiling of microbial communities using 16S rRNA marker gene sequences. Nature Biotechnology 2013;31:814–21.

