## Supplementary figure legend for "Characterization of dynamic age-dependent changes and driver microbes in primate gut microbiota during host’s development and healthy aging via captive crab-eating macaque model"

**Figure S1.** Sparse curve of observed OTUs and sequencing depth.

**Figure S2. Alpha diversity metrics of gut microbiota and their age correlation analysis.** The alpha diversity metrics in the four age groups and their correlation with age are shown including (a,b) Pielou’s evenness, (c, d) phylogenetic diversity, (e, f), observed OTUs; (g,h) Shannon (i,j) and Simpson’s indices were calculated using Spearman correlation. IF, infants; YA, young adults; MA, the middle-aged; EL, the elderly. *: *P*<0.05; **: *P* < 0.01; ***: *P* < 0.001.

**Figure S3. Differential gut microbial phyla among age groups.** Pairwise *P-*values are calculated using nonparametric Kruskal-Wallis test with Tukey post-hoc test. IF, infants; YA, young adults; MA, the middle-aged; EL, the elderly. *: *P*<0.05; **: *P* < 0.01; ***: *P* < 0.001.

**Figure S4. Differential gut microbial genera among age groups.** Pairwise *P-*values are calculated using nonparametric Kruskal-Wallis test with Tukey post-hoc test. IF, infants; YA, young adults; MA, the middle-aged; EL, the elderly. *: *P*<0.05; **: *P* < 0.01; ***: *P* < 0.001.

**Figure S5.** **Spearman** **correlation between differential gut microbial phyla with age.**

**Figure S6.** **Spearman** **correlation between differential gut microbial genera with age.**

**Figure S7. Comparison of additional gut microbial phenotypes predicted by BugBase among the four age groups.** Pairwise *P-*values are calculated using nonparametric Kruskal-Wallis test with Tukey post-hoc test. IF, infants; YA, young adults; MA, the middle-aged; EL, the elderly. *: *P*<0.05; **: *P* < 0.01; ***: *P* < 0.001.

**Figure S8**. **Interactive network between age-associated gut microbial genera and PICRUSt-predicted KEGG modules.** The network is constricted from Spearman correlation between age-associated gut microbial genera and KEGG modules. Correlations with |r| > 0.7 and *P* < 0.05 are shown. Node sizes denote the sum of |r| of the genus from all of its correlations with KEGG modules. Nodes are color green if negatively correlated with age, and yellow if positively correlated with age. Eclipses denote age-associated microbial genera, and diamonds denote age-associated KEGG modules. Genera with their name colored red were driver microbes identified by NetShift analysis. Edges are colored red if r >0, and blue if r < 0.
