## Supplementary figures and images for "Characterization of dynamic age-dependent changes and driver microbes in primate gut microbiota during host’s development and healthy aging via captive crab-eating macaque model"

### Figure S1

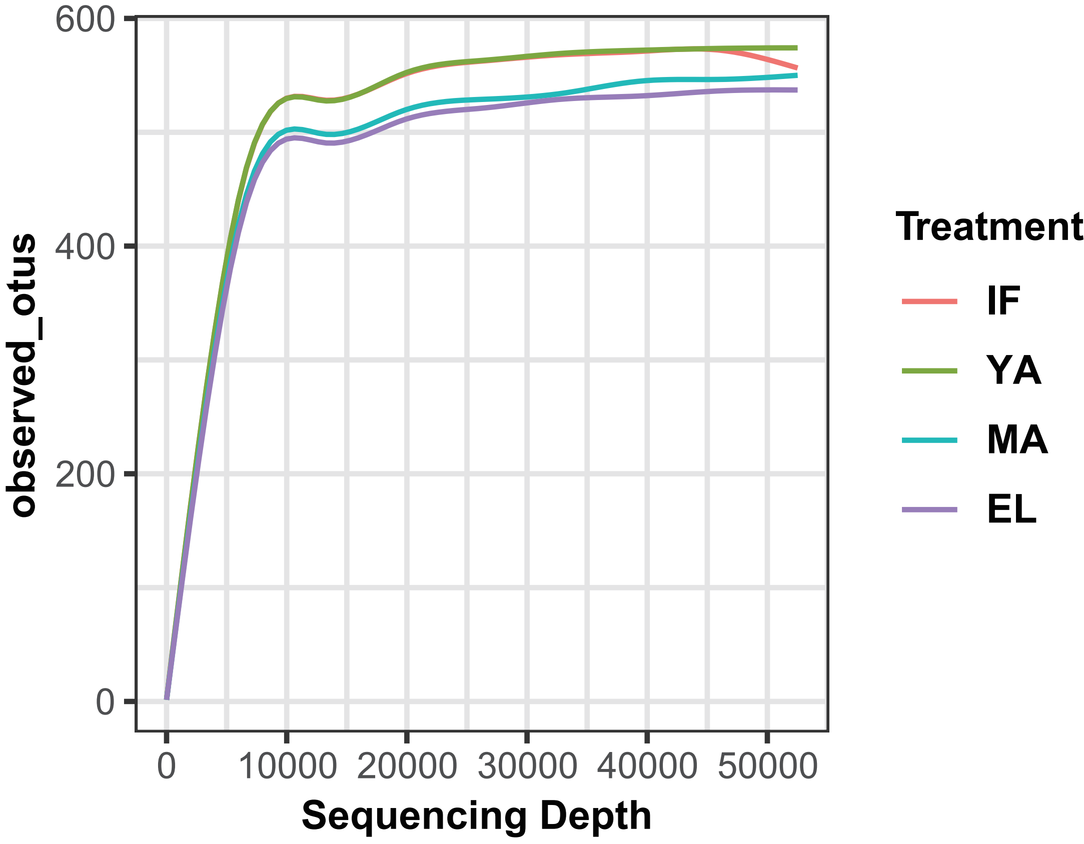

### Figure S2

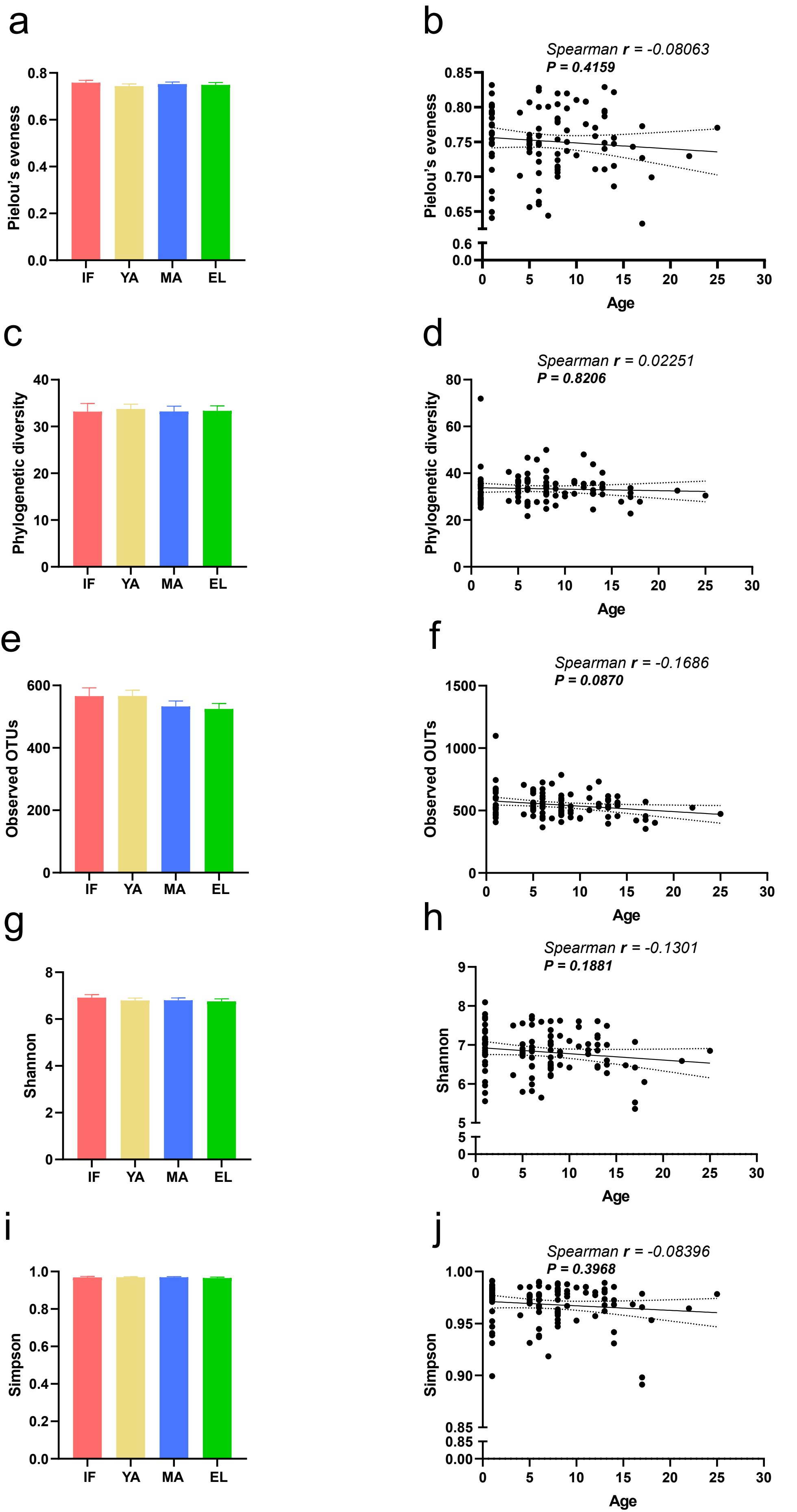

### Figure S3

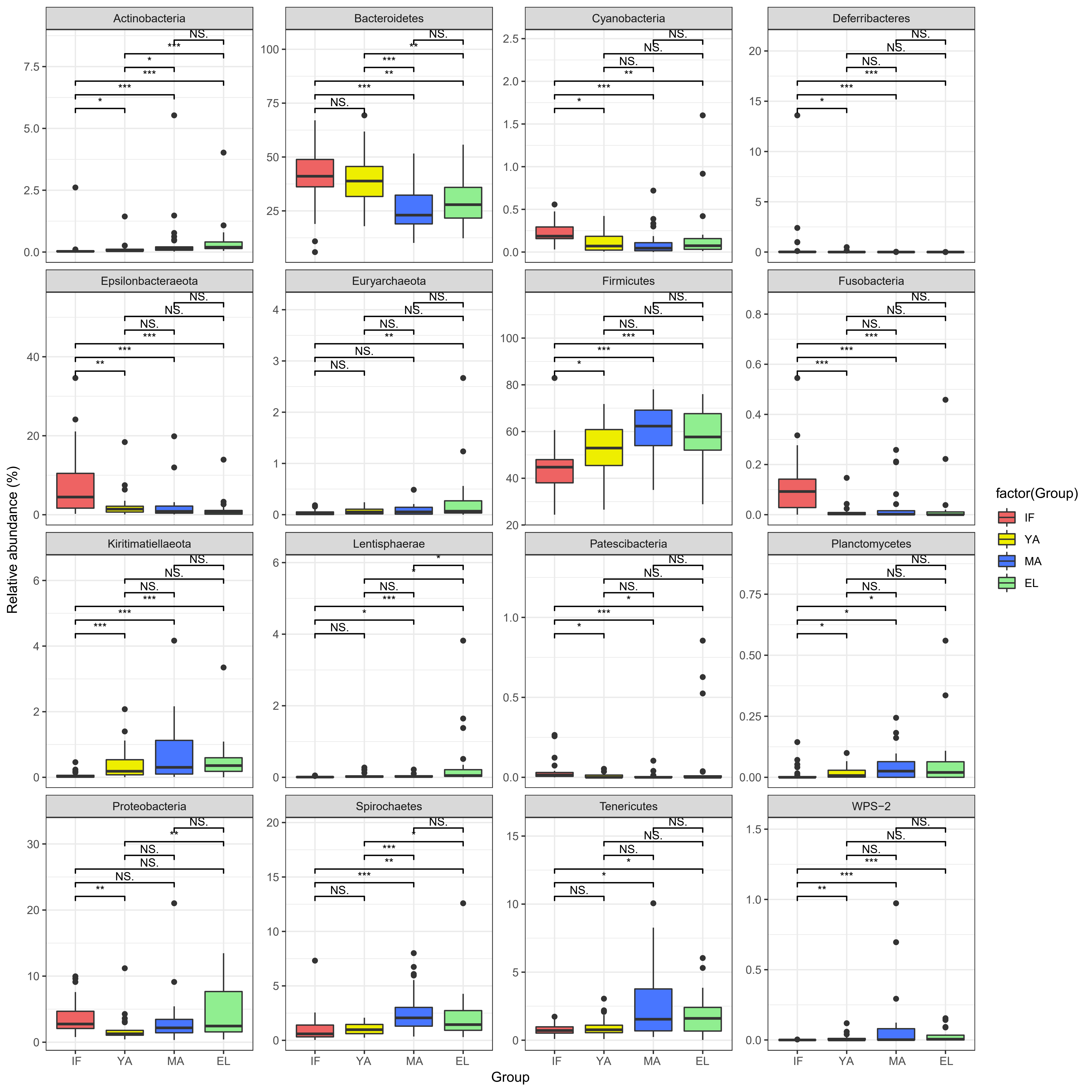

### Figure S4

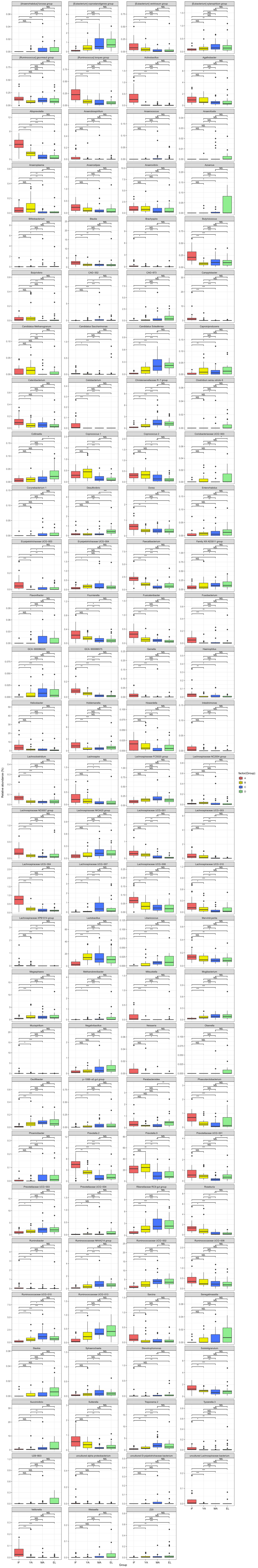

### Figure S5

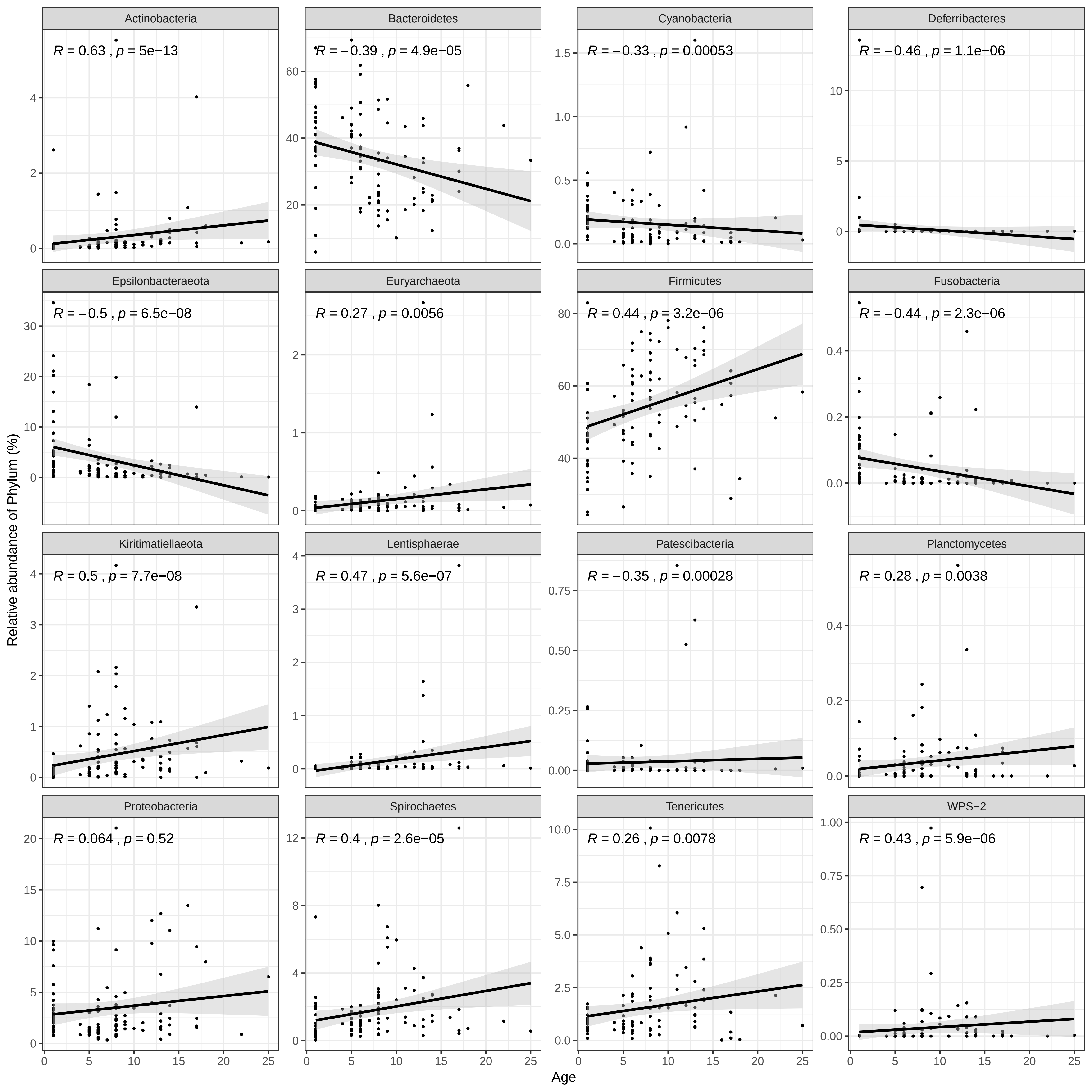

### Figure S6

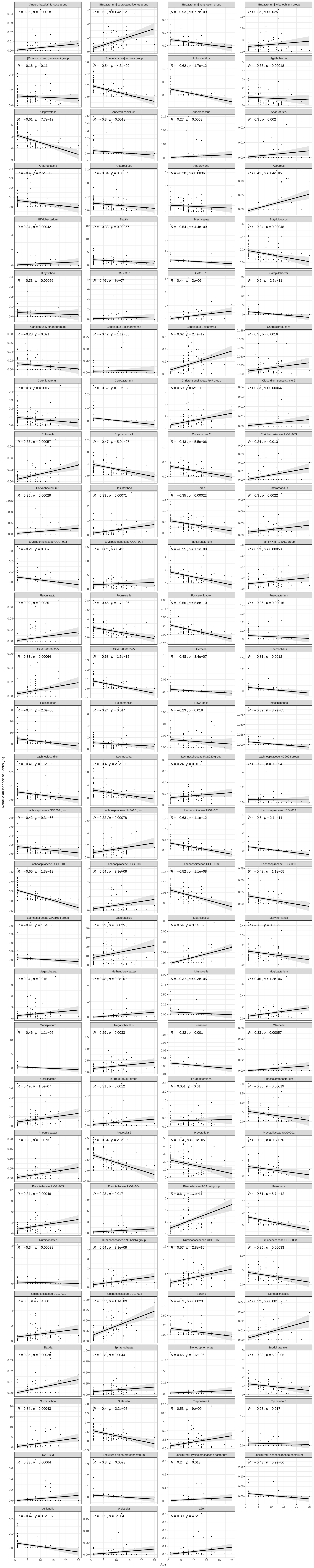

### Figure S7

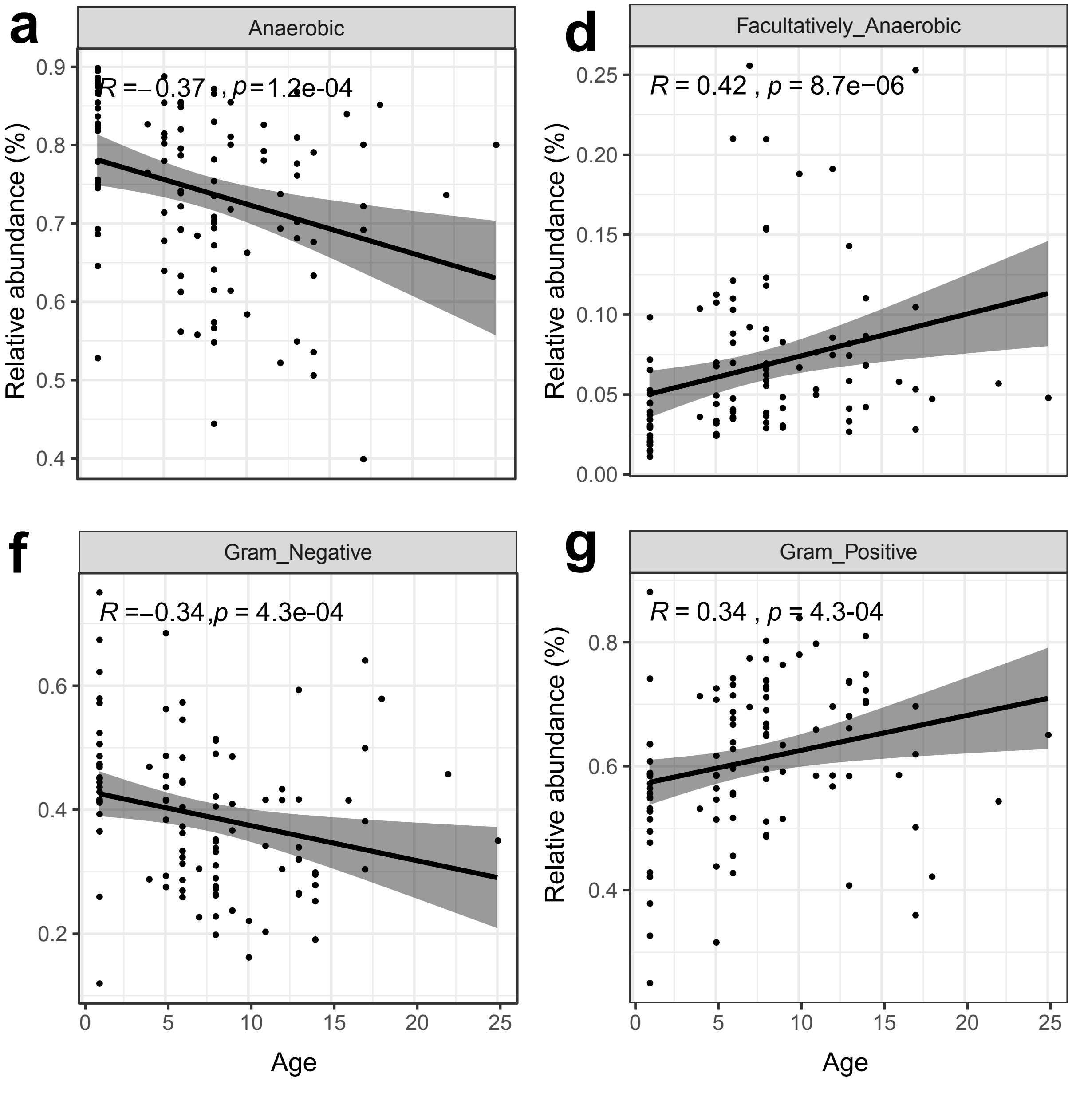

### Figure S8

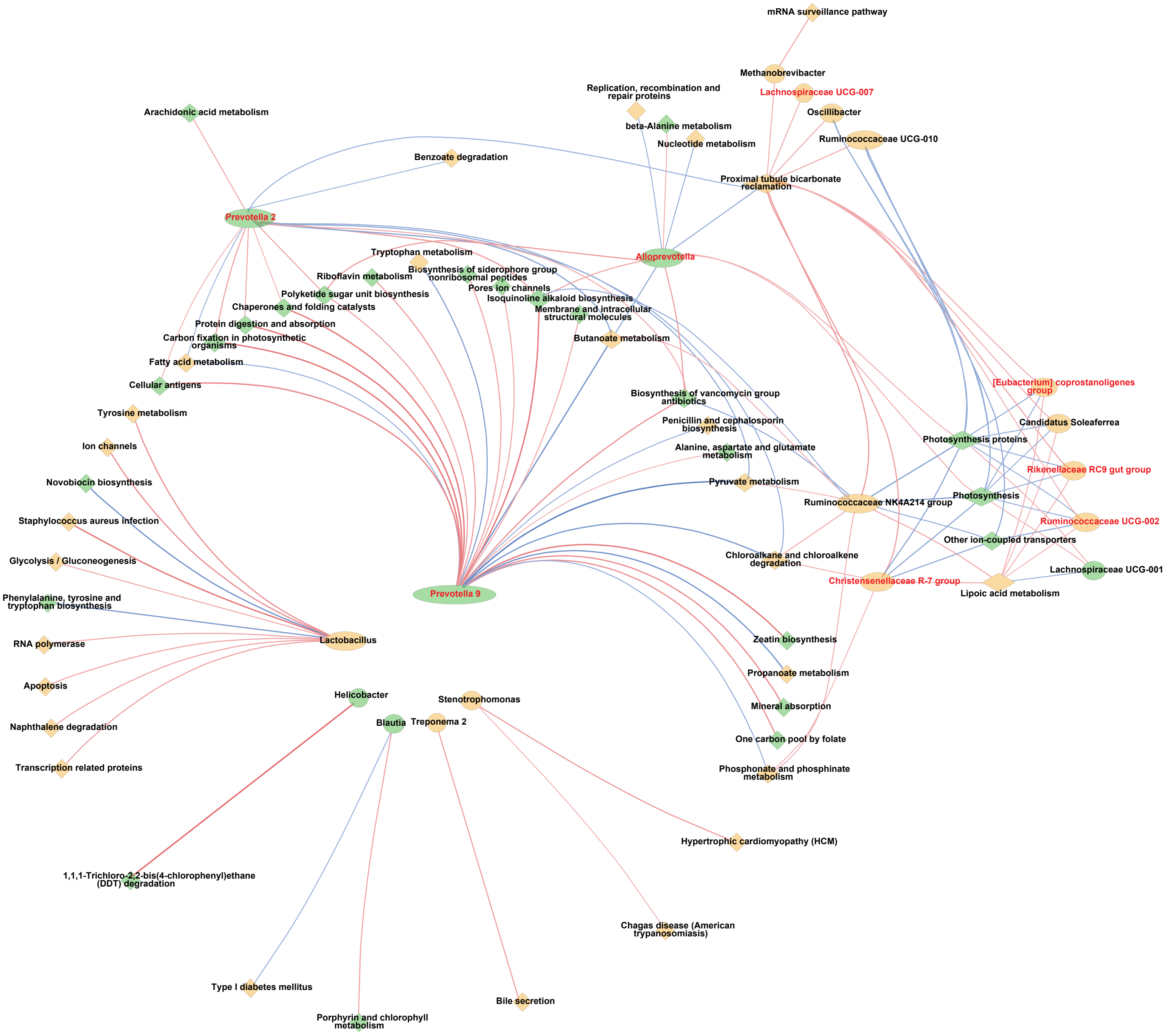
