## Supplementary material for "Characterization of dynamic age-dependent changes and driver microbes in primate gut microbiota during host’s development and healthy aging via captive crab-eating macaque model": Table S1

**Table S1. Sample reads processed with the analysis pipeline in the current study.**

| sample-id | input | filtered | denoised | non-chimeric |
|---|---|---|---|---|
| MAFBL1 | 80078 | 56538 | 56538 | 56104 |
| MAFBL10 | 80046 | 55435 | 55435 | 55028 |
| MAFBL100 | 80178 | 56829 | 56829 | 56381 |
| MAFBL101 | 80087 | 51950 | 51950 | 51666 |
| MAFBL102 | 80144 | 55669 | 55669 | 55202 |
| MAFBL103 | 80248 | 45009 | 45009 | 44752 |
| MAFBL104 | 59009 | 32316 | 32316 | 32042 |
| MAFBL11 | 80138 | 53828 | 53828 | 53618 |
| MAFBL12 | 80083 | 55775 | 55775 | 55440 |
| MAFBL13 | 74503 | 48213 | 48213 | 47314 |
| MAFBL14 | 80020 | 55490 | 55490 | 55308 |
| MAFBL15 | 80111 | 55550 | 55550 | 55145 |
| MAFBL16 | 80100 | 55266 | 55266 | 54794 |
| MAFBL17 | 80141 | 54910 | 54910 | 54560 |
| MAFBL18 | 80133 | 55096 | 55096 | 54927 |
| MAFBL2 | 80169 | 54239 | 54239 | 53657 |
| MAFBL19 | 80182 | 55343 | 55343 | 55031 |
| MAFBL20 | 80119 | 56291 | 56291 | 56291 |
| MAFBL21 | 80149 | 55414 | 55414 | 55287 |
| MAFBL22 | 80202 | 56507 | 56507 | 55853 |
| MAFBL23 | 80228 | 55314 | 55314 | 54707 |
| MAFBL24 | 80081 | 52403 | 52403 | 52066 |
| MAFBL25 | 80099 | 56785 | 56785 | 56545 |
| MAFBL26 | 80052 | 55544 | 55544 | 55151 |
| MAFBL27 | 80051 | 53598 | 53598 | 53598 |
| MAFBL28 | 80079 | 54094 | 54094 | 53696 |
| MAFBL3 | 80078 | 55306 | 55306 | 54891 |
| MAFBL29 | 80063 | 56449 | 56449 | 54954 |
| MAFBL30 | 80126 | 55276 | 55276 | 55096 |
| MAFBL31 | 80166 | 56095 | 56095 | 55477 |
| MAFBL32 | 80213 | 54504 | 54504 | 54152 |
| MAFBL33 | 80047 | 54775 | 54775 | 54264 |
| MAFBL34 | 80117 | 55211 | 55211 | 55207 |
| MAFBL35 | 80130 | 53834 | 53834 | 53666 |
| MAFBL36 | 80037 | 55140 | 55140 | 54916 |
| MAFBL37 | 80156 | 57057 | 57057 | 56818 |
| MAFBL38 | 80311 | 54293 | 54293 | 54153 |
| MAFBL4 | 80120 | 55334 | 55334 | 54942 |
| MAFBL39 | 80093 | 56674 | 56674 | 56624 |
| MAFBL40 | 80028 | 55064 | 55064 | 54971 |
| MAFBL41 | 80171 | 56613 | 56613 | 56511 |
| MAFBL42 | 80101 | 58781 | 58781 | 58432 |
| MAFBL43 | 80175 | 55997 | 55997 | 55701 |
| MAFBL44 | 80125 | 56288 | 56288 | 55889 |
| MAFBL45 | 80171 | 56463 | 56463 | 56117 |
| MAFBL46 | 80210 | 56713 | 56713 | 56402 |
| MAFBL47 | 80188 | 57101 | 57101 | 56851 |
| MAFBL48 | 80125 | 55029 | 55029 | 54799 |
| MAFBL5 | 80112 | 56138 | 56138 | 56108 |
| MAFBL49 | 80188 | 55547 | 55547 | 55147 |
| MAFBL50 | 80159 | 55698 | 55698 | 55394 |
| MAFBL51 | 80096 | 56310 | 56310 | 56180 |
| MAFBL53 | 48642 | 35514 | 35514 | 34270 |
| MAFBL54 | 80226 | 56867 | 56867 | 56252 |

| | | | | |
|---|---|---|---|---|
| MAFBL55 | 80051 | 53742 | 53742 | 53168 |
| MAFBL56 | 80155 | 55715 | 55715 | 55230 |
| MAFBL57 | 80323 | 53880 | 53880 | 53799 |
| MAFBL52 | 81345 | 54549 | 54549 | 54037 |
| MAFBL58 | 80200 | 58022 | 58022 | 57766 |
| MAFBL6 | 80173 | 55124 | 55124 | 54971 |
| MAFBL59 | 80082 | 53800 | 53800 | 53641 |
| MAFBL60 | 80093 | 55199 | 55199 | 55031 |
| MAFBL61 | 80136 | 55843 | 55843 | 55771 |
| MAFBL62 | 80114 | 57319 | 57319 | 56821 |
| MAFBL63 | 80179 | 55607 | 55607 | 55379 |
| MAFBL64 | 80050 | 55215 | 55215 | 54493 |
| MAFBL65 | 80182 | 56630 | 56630 | 56488 |
| MAFBL66 | 80210 | 54781 | 54781 | 54510 |
| MAFBL67 | 80153 | 55414 | 55414 | 55122 |
| MAFBL7 | 80047 | 52818 | 52818 | 52472 |
| MAFBL68 | 80097 | 56161 | 56161 | 55966 |
| MAFBL69 | 80070 | 58009 | 58009 | 57983 |
| MAFBL70 | 80136 | 55337 | 55337 | 55299 |
| MAFBL71 | 80132 | 55872 | 55872 | 55817 |
| MAFBL72 | 80234 | 55784 | 55784 | 55663 |
| MAFBL73 | 80108 | 54567 | 54567 | 54207 |
| MAFBL74 | 80091 | 56484 | 56484 | 56339 |
| MAFBL75 | 80039 | 58664 | 58664 | 57968 |
| MAFBL76 | 85119 | 48053 | 48053 | 47956 |
| MAFBL77 | 84998 | 49572 | 49572 | 49161 |
| MAFBL78 | 65496 | 38456 | 38456 | 38318 |
| MAFBL79 | 80123 | 54125 | 54125 | 53844 |
| MAFBL80 | 80160 | 53529 | 53529 | 53196 |
| MAFBL8 | 80080 | 54718 | 54718 | 54483 |
| MAFBL81 | 86432 | 60742 | 60742 | 59532 |
| MAFBL82 | 80163 | 54980 | 54980 | 54815 |
| MAFBL83 | 80076 | 57469 | 57469 | 57192 |
| MAFBL84 | 80170 | 54922 | 54922 | 54607 |
| MAFBL85 | 80130 | 55340 | 55340 | 55146 |
| MAFBL86 | 80104 | 58483 | 58483 | 58379 |
| MAFBL87 | 80166 | 56867 | 56867 | 56737 |
| MAFBL88 | 80210 | 54714 | 54714 | 54714 |
| MAFBL89 | 80216 | 55048 | 55048 | 54919 |
| MAFBL9 | 80087 | 54890 | 54890 | 54770 |
| MAFBL90 | 80069 | 56094 | 56094 | 56077 |
| MAFBL91 | 80096 | 55027 | 55027 | 54944 |
| MAFBL92 | 80025 | 57025 | 57025 | 56700 |
| MAFBL93 | 80096 | 54211 | 54211 | 53914 |
| MAFBL94 | 80141 | 56705 | 56705 | 55753 |
| MAFBL95 | 80086 | 56578 | 56578 | 56527 |
| MAFBL96 | 80199 | 55551 | 55551 | 55234 |
| MAFBL97 | 80106 | 56215 | 56215 | 56082 |
| MAFBL98 | 80034 | 56660 | 56660 | 56525 |
| MAFBL99 | 80039 | 55310 | 55310 | 54974 |
