## Supplementary material for "Characterization of dynamic age-dependent changes and driver microbes in primate gut microbiota during host’s development and healthy aging via captive crab-eating macaque model": Table S2

**Table S2. Age correlation of KEGG modules.**

| NO. | KEGG module | Spearman correlation coefficient | *P* |
|---|---|---|---|
| 1 | Replication, recombination and repair proteins | 0.640357552 | 2.49E-13 |
| 2 | Lipoic acid metabolism | 0.610067197 | 6.22E-12 |
| 3 | African trypanosomiasis | 0.590235383 | 4.28E-11 |
| 4 | Phosphatidylinositol signaling system | 0.569674657 | 2.77E-10 |
| 5 | Limonene and pinene degradation | 0.567244949 | 3.43E-10 |
| 6 | Photosynthesis proteins | -0.555209797 | 9.56E-10 |
| 7 | Photosynthesis | -0.545199402 | 2.18E-09 |
| 8 | Proximal tubule bicarbonate reclamation | 0.537926477 | 3.89E-09 |
| 9 | Bile secretion | 0.53158224 | 6.39E-09 |
| 10 | beta-Alanine metabolism | -0.520616159 | 1.47E-08 |
| 11 | Penicillin and cephalosporin biosynthesis | 0.518110861 | 1.77E-08 |
| 12 | Naphthalene degradation | 0.517068786 | 1.91E-08 |
| 13 | Glycolysis / Gluconeogenesis | 0.514271923 | 2.35E-08 |
| 14 | Sulfur metabolism | -0.51312726 | 2.55E-08 |
| 15 | Inositol phosphate metabolism | 0.50515242 | 4.53E-08 |
| 16 | Nucleotide metabolism | 0.502555332 | 5.45E-08 |
| 17 | 1,1,1-Trichloro-2,2-bis(4-chlorophenyl)ethane (DDT) degradation | -0.501189297 | 6.00E-08 |
| 18 | Chloroalkane and chloroalkene degradation | 0.491891616 | 1.14E-07 |
| 19 | Phosphonate and phosphinate metabolism | 0.490973726 | 1.21E-07 |
| 20 | Plant-pathogen interaction | -0.48188122 | 2.23E-07 |
| 21 | Butanoate metabolism | 0.478895379 | 2.71E-07 |
| 22 | Oxidative phosphorylation | -0.463339851 | 7.31E-07 |
| 23 | Caprolactam degradation | 0.463021289 | 7.45E-07 |
| 24 | NOD-like receptor signaling pathway | -0.462843111 | 7.54E-07 |
| 25 | Chagas disease (American trypanosomiasis) | 0.457308659 | 1.06E-06 |
| 26 | Biosynthesis of vancomycin group antibiotics | -0.450753965 | 1.57E-06 |
| 27 | Vibrio cholerae pathogenic cycle | -0.450316618 | 1.62E-06 |
| 28 | Progesterone-mediated oocyte maturation | -0.449620102 | 1.68E-06 |
| 29 | Antigen processing and presentation | -0.449620102 | 1.68E-06 |
| 30 | Prostate cancer | -0.448426845 | 1.81E-06 |
| 31 | Benzoate degradation | 0.440678777 | 2.85E-06 |
| 32 | Fatty acid metabolism | 0.43583556 | 3.76E-06 |
| 33 | Selenocompound metabolism | -0.435770768 | 3.77E-06 |
| 34 | Porphyrin and chlorophyll metabolism | -0.433108888 | 4.39E-06 |
| 35 | Apoptosis | 0.422882519 | 7.74E-06 |
| 36 | RNA polymerase | 0.421916035 | 8.16E-06 |
| 37 | Base excision repair | 0.421143928 | 8.51E-06 |
| 38 | Xylene degradation | 0.418325467 | 9.92E-06 |
| 39 | Polyketide sugar unit biosynthesis | -0.416646269 | 1.09E-05 |
| 40 | N-Glycan biosynthesis | -0.413266276 | 1.30E-05 |
| 41 | Protein folding and associated processing | -0.411030945 | 1.46E-05 |
| 42 | Dioxin degradation | 0.404135974 | 2.09E-05 |
| 43 | Riboflavin metabolism | -0.399292757 | 2.68E-05 |
| 44 | Ascorbate and aldarate metabolism | 0.396414903 | 3.10E-05 |
| 45 | Glycerolipid metabolism | 0.395934361 | 3.18E-05 |
| 46 | Isoquinoline alkaloid biosynthesis | -0.395043468 | 3.32E-05 |
| 47 | Nitrogen metabolism | -0.395016472 | 3.33E-05 |
| 48 | Cardiac muscle contraction | -0.393682832 | 3.56E-05 |
| 49 | Staphylococcus aureus infection | 0.393672033 | 3.56E-05 |
| 50 | Meiosis - yeast | 0.392808137 | 3.72E-05 |
| 51 | Ethylbenzene degradation | 0.382803141 | 6.06E-05 |
| 52 | Butirosin and neomycin biosynthesis | 0.380756787 | 6.68E-05 |
| 53 | Influenza A | 0.380574224 | 6.74E-05 |
| 54 | Transcription related proteins | 0.378235291 | 7.53E-05 |

| | | | |
|---|---|---|---|
| 55 | Type I diabetes mellitus | 0.377009638 | 7.98E-05 |
| 56 | Parkinson_s disease | -0.374693317 | 8.90E-05 |
| 57 | Hepatitis C | 0.374660812 | 8.92E-05 |
| 58 | Measles | 0.374660812 | 8.92E-05 |
| 59 | mTOR signaling pathway | 0.374660812 | 8.92E-05 |
| 60 | Cell cycle | 0.374660812 | 8.92E-05 |
| 61 | Other ion-coupled transporters | -0.374390953 | 9.03E-05 |
| 62 | Isoflavonoid biosynthesis | 0.373030046 | 9.62E-05 |
| 63 | Various types of N-glycan biosynthesis | -0.372225814 | 9.99E-05 |
| 64 | Restriction enzyme | 0.371275528 | 0.0001044 |
| 65 | Signal transduction mechanisms | 0.369385755 | 0.0001139 |
| 66 | Carbon fixation in photosynthetic organisms | -0.367220616 | 0.0001258 |
| 67 | Alzheimer_s disease | -0.367085632 | 0.0001266 |
| 68 | Ion channels | 0.363624649 | 0.0001481 |
| 69 | Proteasome | -0.363522061 | 0.0001488 |
| 70 | Tyrosine metabolism | 0.361826665 | 0.0001606 |
| 71 | Cell division | -0.360314847 | 0.0001719 |
| 72 | mRNA surveillance pathway | 0.359801535 | 0.0001759 |
| 73 | Glycine, serine and threonine metabolism | -0.355995367 | 0.0002082 |
| 74 | Prion diseases | -0.351962052 | 0.0002484 |
| 75 | Linoleic acid metabolism | 0.351162948 | 0.0002572 |
| 76 | Chaperones and folding catalysts | -0.350671607 | 0.0002627 |
| 77 | Ribosome Biogenesis | 0.350623013 | 0.0002633 |
| 78 | Bacterial toxins | 0.350509627 | 0.0002646 |
| 79 | Phagosome | 0.349221332 | 0.0002797 |
| 80 | Basal transcription factors | 0.34873864 | 0.0002856 |
| 81 | Pyruvate metabolism | 0.344278776 | 0.0003455 |
| 82 | Lipopolysaccharide biosynthesis | -0.342189228 | 0.0003773 |
| 83 | Metabolism of xenobiotics by cytochrome P450 | 0.335645215 | 0.0004955 |
| 84 | Metabolism of cofactors and vitamins | -0.335326653 | 0.0005021 |
| 85 | Aminobenzoate degradation | 0.334343972 | 0.0005227 |
| 86 | Folate biosynthesis | -0.334333173 | 0.000523 |
| 87 | Epithelial cell signaling in Helicobacter pylori infection | -0.333517871 | 0.0005407 |
| 88 | Bisphenol degradation | 0.331325735 | 0.0005912 |
| 89 | Pores ion channels | -0.328798839 | 0.0006548 |
| 90 | Lipid biosynthesis proteins | 0.324824917 | 0.0007675 |
| 91 | Hypertrophic cardiomyopathy (HCM) | 0.323841365 | 0.000798 |
| 92 | Membrane and intracellular structural molecules | -0.323588466 | 0.000806 |
| 93 | MAPK signaling pathway - yeast | 0.322190034 | 0.0008518 |
| 94 | Phosphotransferase system (PTS) | 0.319771125 | 0.0009364 |
| 95 | Ribosome biogenesis in eukaryotes | 0.319414768 | 0.0009496 |
| 96 | Type II diabetes mellitus | 0.317994739 | 0.0010035 |
| 97 | Vibrio cholerae infection | 0.317640516 | 0.0010173 |
| 98 | Carbon fixation pathways in prokaryotes | -0.311396733 | 0.0012925 |
| 99 | Drug metabolism - cytochrome P450 | 0.309215395 | 0.0014035 |
| 100 | Galactose metabolism | 0.306953067 | 0.0015277 |
| 101 | Lipopolysaccharide biosynthesis proteins | -0.299577555 | 0.0020052 |
| 102 | Function unknown | 0.296019383 | 0.0022807 |
| 103 | alpha-Linolenic acid metabolism | -0.293708461 | 0.0024774 |
| 104 | Novobiocin biosynthesis | -0.292806769 | 0.0025582 |
| 105 | Cellular antigens | -0.290409458 | 0.0027848 |
| 106 | Glyoxylate and dicarboxylate metabolism | -0.287407419 | 0.0030939 |
| 107 | Biosynthesis of unsaturated fatty acids | 0.286462533 | 0.0031974 |
| 108 | Biosynthesis of siderophore group nonribosomal peptides | -0.282056663 | 0.0037221 |
| 109 | Pentose and glucuronate interconversions | 0.280188488 | 0.003967 |
| 110 | Valine, leucine and isoleucine degradation | 0.27991852 | 0.0040035 |
| 111 | Tryptophan metabolism | 0.278984433 | 0.0041323 |
| 112 | Ubiquitin system | 0.274459777 | 0.0048102 |

| | | | |
|---|---|---|---|
| 113 | Pentose phosphate pathway | 0.273514891 | 0.0049636 |
| 114 | Phenylalanine, tyrosine and tryptophan biosynthesis | -0.272877767 | 0.0050696 |
| 115 | Amoebiasis | 0.272003073 | 0.0052183 |
| 116 | DNA replication | 0.270259082 | 0.0055264 |
| 117 | Protein digestion and absorption | -0.269049628 | 0.0057495 |
| 118 | Pathways in cancer | -0.26836931 | 0.0058785 |
| 119 | RNA transport | 0.268039949 | 0.0059418 |
| 120 | Bladder cancer | 0.266364162 | 0.0062737 |
| 121 | Arachidonic acid metabolism | -0.266155576 | 0.0063161 |
| 122 | Flavonoid biosynthesis | 0.265837014 | 0.0063814 |
| 123 | Others | 0.264686953 | 0.0066222 |
| 124 | Amino acid metabolism | 0.26080482 | 0.0074953 |
| 125 | Inorganic ion transport and metabolism | -0.257581408 | 0.008296 |
| 126 | D-Alanine metabolism | 0.256415148 | 0.0086038 |
| 127 | Tropane, piperidine and pyridine alkaloid biosynthesis | -0.251237171 | 0.0100949 |
| 128 | Pantothenate and CoA biosynthesis | -0.247727593 | 0.0112302 |
| 129 | Chlorocyclohexane and chlorobenzene degradation | -0.242414633 | 0.0131613 |
| 130 | Homologous recombination | 0.240254892 | 0.0140254 |
| 131 | Flagellar assembly | -0.234720558 | 0.0164684 |
| 132 | Transcription factors | 0.234094234 | 0.0167668 |
| 133 | Cyanoamino acid metabolism | -0.231389159 | 0.0181098 |
| 134 | Zeatin biosynthesis | -0.224229621 | 0.0221209 |
| 135 | Bacterial invasion of epithelial cells | 0.224159429 | 0.0221637 |
| 136 | Alanine, aspartate and glutamate metabolism | -0.223959653 | 0.022286 |
| 137 | Protein processing in endoplasmic reticulum | -0.222383043 | 0.0232712 |
| 138 | Hematopoietic cell lineage | 0.221140906 | 0.0240735 |
| 139 | Pertussis | 0.220131514 | 0.0247427 |
| 140 | Taurine and hypotaurine metabolism | 0.218484712 | 0.0258686 |
| 141 | Ubiquinone and other terpenoid-quinone biosynthesis | -0.217550624 | 0.0265264 |
| 142 | Propanoate metabolism | 0.217464235 | 0.0265879 |
| 143 | Glycosphingolipid biosynthesis - lacto and neolacto series | 0.211130538 | 0.0314439 |
| 144 | Mineral absorption | -0.206557547 | 0.0353988 |
| 145 | Flavone and flavonol biosynthesis | 0.204613781 | 0.0372023 |
