## Supplementary material for "Characterization of dynamic age-dependent changes and driver microbes in primate gut microbiota during host’s development and healthy aging via captive crab-eating macaque model": Table S3

**Table S3. Network parameters caculated by NetShift.**

| S_ID | n(control) | n(case) | core(case) | Union | Intersect | Exclusive | Jaccard-score | NESH-score | DelBet | COM |
|---|---|---|---|---|---|---|---|---|---|---|
| YA vs. IF | | | | | | | | | | |
| Prevotellaceae UCG-003 | 1 | 3 | 3 | 3 | 1 | 2 | 0.333 | 1.833 | 0 | 1 |
| Helicobacter | 1 | 1 | 1 | 2 | 0 | 1 | 0 | 1.75 | 0 | 2 |
| Lactobacillus | 1 | 1 | 1 | 2 | 0 | 1 | 0 | 1.75 | 0 | 2 |
| Ruminococcaceae UCG-014 | 1 | 1 | 1 | 2 | 0 | 1 | 0 | 1.75 | 0 | 1 |
| **Prevotella 9** | 2 | 4 | 3 | 4 | 2 | 2 | 0.5 | 1.5 | 1 | 1 |
| Alloprevotella | 2 | 3 | 3 | 3 | 2 | 1 | 0.667 | 0.917 | 0 | 1 |
| Prevotella 2 | 3 | 3 | 3 | 3 | 3 | 0 | 1 | 0 | -1 | 1 |
| MA vs. YA | | | | | | | | | | |
| **Dialister** | 1 | 5 | 4 | 5 | 1 | 4 | 0.2 | 1.933 | 0.038 | 2 |
| **Megasphaera** | 1 | 5 | 4 | 5 | 1 | 4 | 0.2 | 1.933 | 0.015 | 2 |
| **Christensenellaceae R-7 group** | 3 | 6 | 4 | 8 | 1 | 5 | 0.125 | 1.917 | 0.104 | 1 |
| Ruminococcaceae UCG-010 | 2 | 5 | 4 | 6 | 1 | 4 | 0.167 | 1.833 | -0.129 | 1 |
| **Rikenellaceae RC9 gut group** | 4 | 7 | 4 | 9 | 2 | 5 | 0.222 | 1.75 | 0.103 | 2 |
| Ruminococcaceae UCG-014 | 1 | 1 | 1 | 2 | 0 | 1 | 0 | 1.583 | 0 | 1 |
| **[Eubacterium] coprostanoligenes group** | 3 | 5 | 4 | 6 | 2 | 3 | 0.333 | 1.417 | 0.009 | 1 |
| **Ruminococcaceae UCG-005** | 3 | 5 | 4 | 6 | 2 | 3 | 0.333 | 1.417 | 0.015 | 2 |
| **Ruminococcaceae UCG-002** | 7 | 10 | 4 | 11 | 6 | 4 | 0.545 | 1.152 | 0.636 | 1 |
| Anaerovibrio | 1 | 2 | 2 | 2 | 1 | 1 | 0.5 | 1.083 | 0 | 1 |
| Prevotella 2 | 7 | 5 | 4 | 9 | 3 | 2 | 0.333 | 1.056 | -0.019 | 3 |
| Lactobacillus | 4 | 1 | 1 | 4 | 1 | 0 | 0.25 | 0.75 | -0.414 | 4 |
| Prevotella 9 | 13 | 12 | 4 | 15 | 10 | 2 | 0.667 | 0.633 | 0 | 3 |
| Alloprevotella | 6 | 4 | 3 | 6 | 4 | 0 | 0.667 | 0.333 | 0.012 | 3 |
| Agathobacter | 2 | 2 | 2 | 2 | 2 | 0 | 1 | 0 | 0 | 3 |
| Succinivibrio | 1 | 1 | 1 | 1 | 1 | 0 | 1 | 0 | 0 | 4 |
| EL vs. MA | | | | | | | | | | |
| **Ruminococcaceae UCG-014** | 2 | 4 | 2 | 6 | 0 | 4 | 0 | 2.067 | 0.23 | 1 |
| **Holdemanella** | 1 | 2 | 1 | 3 | 0 | 2 | 0 | 1.867 | 0.24 | 1 |
| Faecalibacterium | 1 | 1 | 1 | 2 | 0 | 1 | 0 | 1.6 | 0 | 3 |
| Treponema 2 | 9 | 3 | 2 | 12 | 0 | 3 | 0 | 1.55 | -0.194 | 4 |
| **Succinivibrio** | 1 | 3 | 2 | 3 | 1 | 2 | 0.333 | 1.533 | 0.06 | 4 |
| **Rikenellaceae RC9 gut group** | 7 | 4 | 2 | 10 | 1 | 3 | 0.1 | 1.5 | 0.309 | 4 |
| Oribacterium | 2 | 1 | 1 | 3 | 0 | 1 | 0 | 1.433 | 0 | 1 |
| **Alloprevotella** | 5 | 5 | 2 | 8 | 2 | 3 | 0.25 | 1.425 | 0.234 | 2 |
| **Lachnospiraceae UCG-007** | 3 | 3 | 2 | 5 | 1 | 2 | 0.2 | 1.4 | 0.124 | 1 |
| **Prevotella 2** | 7 | 3 | 2 | 9 | 1 | 2 | 0.111 | 1.311 | 0.054 | 3 |
| Ruminococcus 1 | 6 | 1 | 1 | 7 | 0 | 1 | 0 | 1.243 | -0.006 | 1 |
| Christensenellaceae R-7 group | 8 | 2 | 2 | 9 | 1 | 1 | 0.111 | 1.1 | -0.103 | 2 |
| Lactobacillus | 1 | 2 | 2 | 2 | 1 | 1 | 0.5 | 1.1 | 0 | 4 |
| **Megasphaera** | 7 | 2 | 1 | 8 | 1 | 1 | 0.125 | 1.1 | 0.056 | 1 |
| Ruminococcaceae NK4A214 | 6 | 2 | 2 | 7 | 1 | 1 | 0.143 | 1.1 | -0.006 | 2 |
| Subdoligranulum | 1 | 2 | 2 | 2 | 1 | 1 | 0.5 | 1.1 | 0 | 3 |
| Dialister | 4 | 1 | 1 | 4 | 1 | 0 | 0.25 | 0.75 | -0.007 | 1 |
| Ruminococcaceae UCG-010 | 4 | 1 | 1 | 4 | 1 | 0 | 0.25 | 0.75 | -0.002 | 3 |
| Prevotella 9 | 17 | 10 | 2 | 18 | 9 | 1 | 0.5 | 0.656 | 0 | 3 |
| Ruminococcaceae UCG-002 | 11 | 4 | 2 | 11 | 4 | 0 | 0.364 | 0.636 | -0.258 | 2 |
| Agathobacter | 2 | 1 | 1 | 2 | 1 | 0 | 0.5 | 0.5 | 0 | 3 |
| Anaerovibrio | 2 | 1 | 1 | 2 | 1 | 0 | 0.5 | 0.5 | 0 | 3 |

IF, infants; YA, young adults; MA, the middle-aged; EL, the elderly.

**Red bold face denotes identified drivers.**
